# AAV2 Crosslinks Actin Filaments: Implications for AAV Gene Therapy Vector Design

**DOI:** 10.64898/2026.05.08.723880

**Authors:** Mitchell Gulkis, James B. Heidings, Juha T. Huiskonen, Anjelique Sawh-Gopal, Jane Hsi-Bell, Mark Potter, Xiyong Song, Tarun E. Hutchinson, Antonette Bennett, Mario Mietzsch, Jonathan E. Bird, Robert McKenna

## Abstract

Adeno-associated virus (AAV) capsids are important gene therapy vectors, allowing for the one-time treatment of monogenetic disorders, with durable gene expression lasting for years. However, despite the clinical success of AAV usage, low transduction efficiencies require high dosage to achieve therapeutic efficacy, resulting in prohibitive costs and rare, but life-threatening immune responses. One key knowledge gap is how the capsids traffic to the nucleus following endocytosis. Here, we identify a direct interaction between AAV2 and actin filaments. Our results show that AAV2 bundles multiple actin filaments in a highly periodic hexagonal lattice. Therefore, we propose that AAV2 is the founding member of a new class of actin binding proteins, the actin bundling viruses (ABVs). Using cryogenic electron microscopy, we determined the structure of AAV2-actin filaments, identified the interaction interfaces, and engineered capsid variants that no longer bundle actin. Together, our results suggest that the AAV2-actin interaction may be responsible for trapping capsids and mediating peri-nuclear accumulation. Overall, this interaction expands the current understanding of AAV2 biology and offers new directions for capsid engineering.

## Introduction

Adeno-associated virus (AAV) capsids are important vectors for gene therapy, currently being used in nine FDA-approved treatments and over a hundred ongoing clinical trials.^1–5^ One of the major advantages of AAV-mediated gene therapy is the durable expression of the transgene, which is primarily maintained as an episome in non-dividing cells.^1–5^ Therefore, AAV-mediated gene therapy results in therapeutic efficacy that is measured in years, oftentimes extending the lives of individuals treated as well as their quality of life.^2,3,6^ Despite these advantages and the major benefit to patients, AAV-mediated gene therapy has several key drawbacks.

One major limitation is the requirement for high dose of therapeutic particles for treatment. Patients are often dosed with 10^12^-10^14^ vector genomes per kilogram body weight.^7^ These large doses can result in unwanted side effects, such as triggering hyperactive immune responses, which can be fatal in rare instances.^7^ Additionally, as a viral vector, AAV capsids are significantly more expensive to research, develop, and produce when compared to small molecule therapies, which combined with the large dosage, results in seven-figure price tags for AAV-mediated gene therapies.^8,9^ The high dose of AAV transgene or vector genomes is required because the vast majority of administered vectors never reach the nucleus to express their therapeutic payload. The multiplicity of infection (MOI) of AAV particles is approximately 10^5^, orders of magnitude higher than other viruses, such as adenovirus that has a MOI of 5.^10^ While factors such as pre-existing immunity and binding of AAV to off-target tissue certainly also play a role in the poor transduction efficiency of AAV, a significant limiting factor is our understanding of the route of cellular transduction by the capsid.^11,12^

Prior work has shown that the AAVs follow a multi-step pathway from injection to expression of the therapeutic transgene.^12^ First, the capsid has to navigate the complex environment of the body and bind to the target cell type, mediated mainly by cell-surface exposed receptors and glycans.^13–15^ Once bound, the AAV undergoes receptor-mediated endocytosis and enters the endo-lysosomal pathway. Here, continuous acidification of the endosome results in conformational changes in the capsid that externalizes a phospholipase A2 (PLA2) domain that is normally situated in the interior of the capsid. The traditional belief is that the PLA2 domain degrades the membrane of the endo-lysosome and releases the capsid into the cytoplasm. More recent research suggests that the endosomal AAVs could be trafficked to the trans-Golgi network, where they escape into the cytoplasm.^12^ Regardless of the pathway, once in the cytoplasm AAVs preferentially accumulate in the peri-nuclear region, specifically near the microtubule organizing centre (MTOC).^16^ Subsequently, the capsid can cross the nuclear membrane, possibly through the nuclear pore, and release the therapeutic transgene.

While much work has been done to understand AAV cell surface binding, the externalization of the PLA2 domain, and the escape of the endo-lysosomal pathway, there are still open questions on how the capsids are trafficked to the nucleus. Previous studies have implied a role of microtubules for trafficking AAVs to the MTOC.^16–20^ However, when microtubules are depolymerized by nocodazole prior to infection, transduction still occurs, albeit less efficiently, suggesting that another cytoskeletal element is also important for transduction.^16^ Additionally, when microtubules are depolymerized by nocodazole after peri-nuclear accumulation, the transduction rate increases, likely due to an increase in the amount of AAV capsids that are able to enter the nucleus.^17^ Interestingly, this increase in transduction is dependent on polymerized actin filaments, as well as the RhoA-ROCK signalling pathway, which rearranges actin filaments in response to microtubule depolymerization.^17^ Finally, in addition to microtubules, actin filaments are concentrated within the MTOC.^21–23^ Taken together, these three observations suggest that actin filaments may be important for AAV transduction.

Actin is ubiquitously expressed in all cell types and is critical for cell motility, mechanical stability, and intercellular trafficking.^24–26^ In fact, actin is the most highly abundant intracellular protein.^26^ Cellular protrusions such as stereocilia and filopodia as well as muscle fibers are especially rich in actin. Actin undergoes highly regulated, dynamic polymerization and depolymerization from globular actin monomers (G-actin) to filamentous actin polymers (F-actin). It is well known that many other viruses exploit both direct and indirect interactions with the actin cytoskeleton during viral entry, replication, and egress.^27–31^ Therefore, it would not be unprecedented if AAV also utilized the actin cytoskeleton for cellular trafficking.

In this work, we show that the clinically used adeno-associated virus 2 (AAV2) capsid co-localizes with actin filaments in cells and that AAV2 bundles actin filaments *in vitro*. Total internal reflection fluorescence microscopy (TIRFM) reveals the kinetics of actin filament bundling in real time. Furthermore, by utilizing multiple modes of electron microscopy, including negative stain, cryogenic single particle analysis, and cryogenic tomography, we demonstrate the mechanism of bundling and identify the capsid-actin residues which mediate the interaction. Finally, targeted mutagenesis resulted in the generation of an AAV2 variant that no longer bundles actin filaments. Taken together, our work uncovers a direct interaction between AAV2 and actin filaments, providing a structural model for how AAV2 interacts with one of the cell’s most abundant proteins. Future research into the biological function of this interaction will be critical for understanding AAV biology, and modulation of the interaction interface will offer important avenues to increase the transduction efficiency of gene therapy vectors through structure-aided rational design.

## Results

### AAV2 bundles actin filaments

To investigate whether actin is implicated in the trafficking of AAV2, we first used confocal immunofluorescence to visualize the localization of each in fixed cells (Figure 1A).We observed strong spatial co-localization between AAV2 and actin filaments, however due to the resolution of light microscopy, we could not distinguish whether these capsids were directly associated with actin filaments, the cell membrane, or both. Since AAV2 is an icosahedral virus, made up of 60 copies of the capsid protein, we reasoned that if the interaction was direct, each capsid should be able to bind to multiple actin filaments, resulting in filament crosslinking and bundling into larger macromolecular assemblies. To test for direct binding and crosslinking activity, we employed a centrifugation-based assay that differentially sediments actin filaments from cross-linked, bundled actin filaments at low speed (10,000 x *g*). In control experiments, neither AAV2, actin filaments, nor actin filaments plus bovine serum albumin significantly sedimented during centrifugation and instead remained in the supernatant (Figure 1B, middle). As a positive control, we confirmed that the majority of actin filaments shifted into the pellet fraction in response to artificially induced bundling of actin filaments using 30 mM MgCl_2_ (Figure 1B).^32,33^ When AAV2 and actin filaments were pre-incubated together before centrifugation, AAV2 sedimented almost exclusively in the pellet fraction along with the majority of actin filaments (Figure 1B, right). We conclude that AAV2 can directly bind and crosslink actin filaments without any additional accessory proteins required.

**Figure 1.**
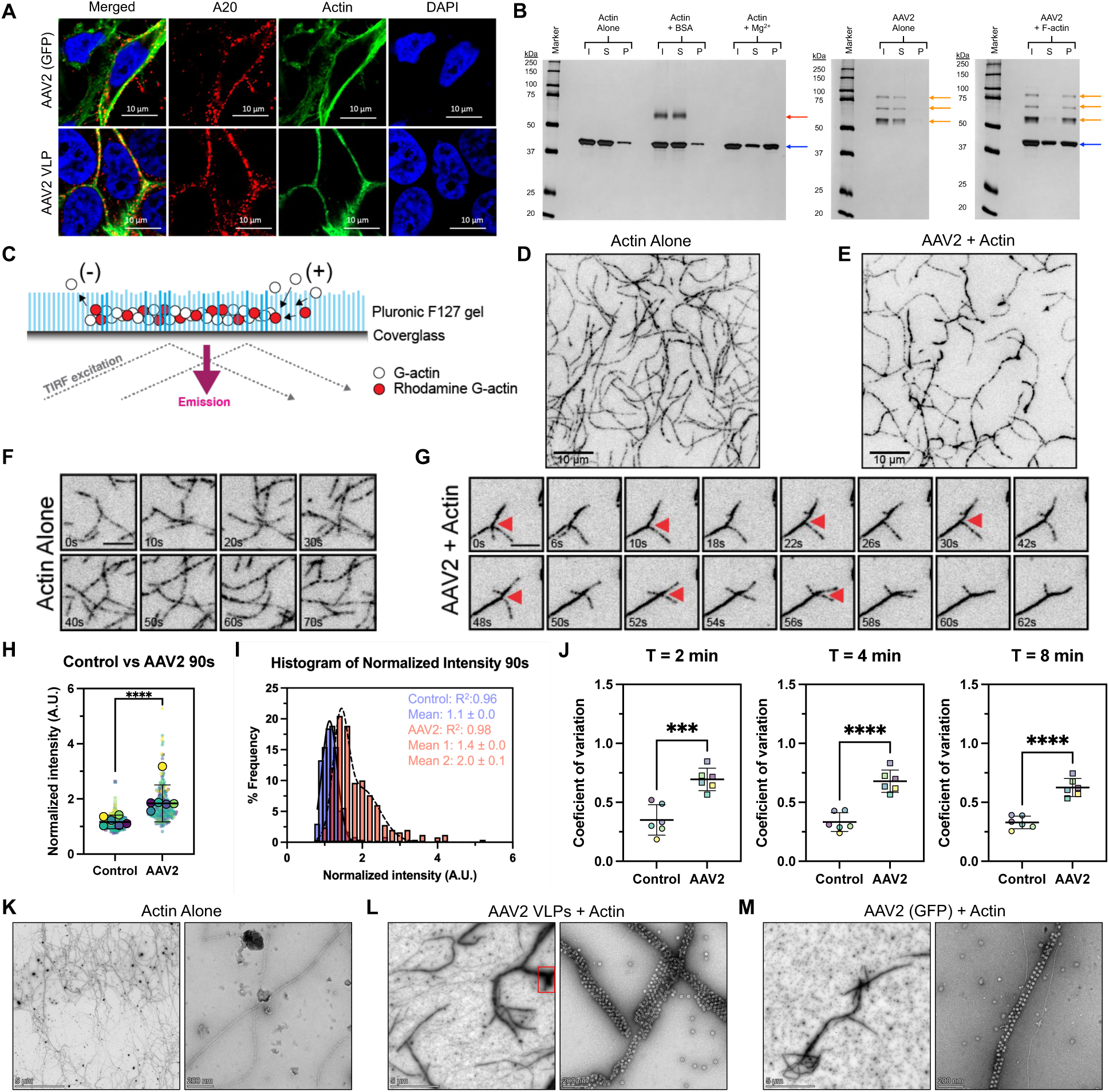
Adeno-associated virus 2 bundles actin filaments. (A) Immunofluorescent confocal microscopy images of HeLa cells infected with either AAV2 packaging a GFP transgene 15 minutes post-infection (top) or AAV2 VLPs 2 hours post-infection (bottom). A20 is an antibody that recognizes intact AAV capsids. (B) Silver stained SDS-PAGE gels containing fractions from a centrifugation-based actin bundling assay. “I” represents the input fraction, “S” represents the supernatant fraction, “P” represents the pellet fraction. The left gel shows the results from the actin alone negative control, the actin + bovine serum albumin negative control, and the actin + Mg^2+^ positive control. The red arrow indicates the BSA band and the blue arrow indicates the actin band. The middle gel shows the results from the AAV2 alone negative control. The orange arrows indicate the AAV2 bands. The right gel shows the results from the AAV2 + actin test reaction. The orange arrows indicate the AAV2 bands and the blue arrow indicates the actin band. (C) Cartoon schematic illustrating the Pluronic F127-based TIRFM actin polymerization and bundling assay. (D-E) Full frame still TIRFM images of actin alone (D) and actin with AAV2 (E) at 10 minutes incubation time. Scale bar is 10 μm. (F-G) Cropped frame series for actin alone (F) and actin with AAV2 (G). Scale bar is 5 μm. Red arrows in (G) denote actin filaments that are bundled over time by AAV2. (H) Normalized pixel intensity for actin alone or actin with AAV2. **** indicates P < 0.0001 using a Mann Whitney T test. (I) Histogram of normalized pixel intensities for actin alone (blue) or actin with AAV2 (orange) fit with either a single Gaussian (actin alone) or a double Gaussian distribution (actin with AAV2). The correlation coefficient, mean, and standard deviation are shown in the upper right corner, colored as above. (J) Coefficient of variation analysis for two minutes of reaction time (left), four minutes of reaction time (middle), and eight minutes of reaction time (right). *** indicates P = 0.0005 using a Welches T test, and **** indicates P < 0.0001 using a Welches T test. (K-M) Negative stain electron micrographs of actin filaments alone (K), actin filaments with AAV2 VLPs (L), or actin filaments with AAV2 packaging a GFP transgene (M). Left images are at 3,400X magnification with a scale bar of 5 μm and right images are at 45,000X magnification with a scale bar of 200 nm.

We next studied the formation of AAV crosslinked actin filaments using time-lapse total internal reflection fluorescence microscopy (TIRFM). Polymerization of G-actin monomers (20% rhodamine coupled actin) was initiated by the addition of 50 mM KCl, 2 mM MgCl_2_, and 0.5 mM EGTA and immediately introduced to the TIRFM flow cell. To facilitate actin crosslinking events, we functionalized the flow cell using a Pluronic F-127 gel to retain actin filaments within the evanescent field whilst allowing for 2D diffusional encounters.^34–36^ In the absence of AAV2, G-actin polymerized into filaments that diffused across the flow cell surface and flexed due to their relatively low (∼ 10 microns) persistence length (Figure 1D,F and Supplementary Video 1-2). Quantification of filament intensities at 90 seconds revealed a single population consistent with these filaments not being crosslinked (Figure 1I). When AAV2 was included in the actin polymerization reaction, an interconnected network of actin filaments assembled that no longer diffused on the flow cell surface, and was rigid indicating a marked increase in persistence length (Figure 1E,G and Supplementary Video 1-2). Using time-lapse TIRFM, we directly visualized bundling events, with individual actin filaments being “zippered” into a larger parent bundle (Figure 1G, red arrow). Filament intensities in the presence of AAV2 were bimodal, with a significant proportion at twice the modal value, indicating the presence of actin bundling (Figure 1I). Analysis of filament fluorescence revealed an increased co-efficient of variation (COV) induced by AAV2 at multiple time points (Figure 1J), similar with other known actin-bundling proteins.^34,37^ Our data demonstrates that AAV2 promotes actin-filament bundling and highlights actin binding as a hitherto unexplored aspect of AAV biology.

To reveal the mechanism of bundling we visualized AAV2 complexed with actin filaments using negative stain electron microscopy. Actin filaments alone appeared as a 7 nm wide double helix, as expected (Figure 1K). Upon addition of AAV2, bundles formed that were microns long and extremely electron dense, and in some regions so dense it became opaque to imaging (Figure 1L, red box). At higher magnifications, AAV2 capsids are clearly visualized bound between co-linear actin filaments (Figure 1L, right). These imaging experiments further validated that each icosahedral capsid has multiple actin binding sites, and likewise each helical filament has multiple AAV2 binding sites. Together, this results in the formation of large AAV2-actin bundles, which we also observed in sedimentation assays and TIRFM assays. We confirmed this interaction with seven independent preparations of AAV2 and three independent preparations of actin across three buffer systems, rigorously indicating the robust bundling activity of AAV2. However, due to the resolution limit of negative stain EM, we could not determine whether each binding interface was identical, or whether there were multiple binding-competent conformations. Additionally, we could not determine the number of bound filaments, or the location of the binding interface. Higher resolution data were needed to obtain this information.

For most of these experiments, virus-like particles (VLPs) lacking a packaged transgene were used. Packaging of the transgene reduces the isoelectric point of the overall capsid but does not change the surface topology of the capsid.^38^ Therefore, if the interaction was only mediated by charge, and was not a specific interaction, it would be possible that packaged capsids would not bind. To explore this possibility, AAV2 capsids packaging a GFP transgene were purified and separated from unpackaged capsids using an iodixanol density gradient then incubated with actin filaments. Negative stain electron microscopy confirms that packaged AAV2 capsids also bind and bundle actin filaments similarly to VLPs, supporting the hypothesis that the interaction between AAV2 and actin filaments is specific (Figure 1M).

While imaging, we noticed that AAV2 capsids appeared to bind actin filaments with striking periodicity (Supplementary Figure 1A-C). When periodic particles were picked with a box size able to accommodate multiple capsids, class averages showed a regular 2D-array of AAV2 capsids. The capsids are periodically spaced approximately 360 Å apart in the direction of the actin filament and approximately 300 Å perpendicular to it (Supplementary Figure 1D-E). These distances correspond well to the 36 nm half helical repeat distance of actin filaments and the distance between two half capsids (25 nm) plus an actin filament (7 nm).^39–41^ The Fourier transform of these 2D-class averages reveals high periodicity with the presence of clear layer lines (Supplementary Figure 1F-G).

We reasoned that binding of AAV2 to actin filaments may be cooperative; once one capsid binds to two filaments, the filaments would be perfectly spaced for further AAV2 binding. To test this, we conducted a time-course experiment, quenching the reaction via addition of the negative stain reagent. When AAV2 and actin filaments were mixed on grid and allowed one minute to adsorb, then immediately stained, micron-scale bundles were not visible, but well-defined local tracks of AAV2 were observed, consistent with cooperativity (Supplementary Figure 2A-B). Interestingly, given one minute longer incubation, micron-scale bundles were readily observed (Supplementary Figure 2C-D) and these bundles grew in size and frequency as a function of time (Supplementary Figure 2C-H). Together, these results suggest that AAV2 binding may be cooperative.

### Structure determination of the AAV2-actin complex

After vitrification of AAV2 incubated with actin filaments, clear bundles were visualized with high periodicity (Supplementary Figure 3A). However, since actin filaments are microns long, they are confined to run approximately parallel to the grid surface, During routine cryo-EM data processing, it became apparent that AAV2 bound in a specific location along the helical repeat of actin filaments, which resulted in a dominant preferred orientation, with only two views of AAV2 bound to actin filaments (Supplementary Figure 3C-D). Furthermore, each capsid was bound to between one to six co-linear actin filaments and the binding sites were heterogenous between each capsid. Finally, due to limited resolution and map anisotropy, we could not rule out the possibility that the polarity of each filament was also heterogenous between capsids. Despite these limitations, we reconstructed the AAV2-actin complex to 5 Å resolution using multiple tilted data sets, particle subtraction, multiple rounds of 3D classification, symmetry relaxation, and per-particle CTF refinement (Supplementary Figure 4, Supplementary Figure 6C and Supplementary Table 1). In this reconstruction, the AAV2 capsid is clearly resolved, with six actin filaments bound with pseudo-6-fold symmetry around the icosahedral 3-fold axis, with each filament binding across the capsid 2-fold symmetry axis, making contacts with two adjacent 3-fold protrusions (Figure 2A and 2C). Two distinct conformations of actin are seen bound to alternating pairs of 3-fold protrusions, which we refer to as Type A and Type B interfaces (Figure 2B – light blue *versus* dark blue). Interestingly, a partial capsid can be seen for the next and previous capsid in the AAV2-actin bundle.

**Figure 2.**
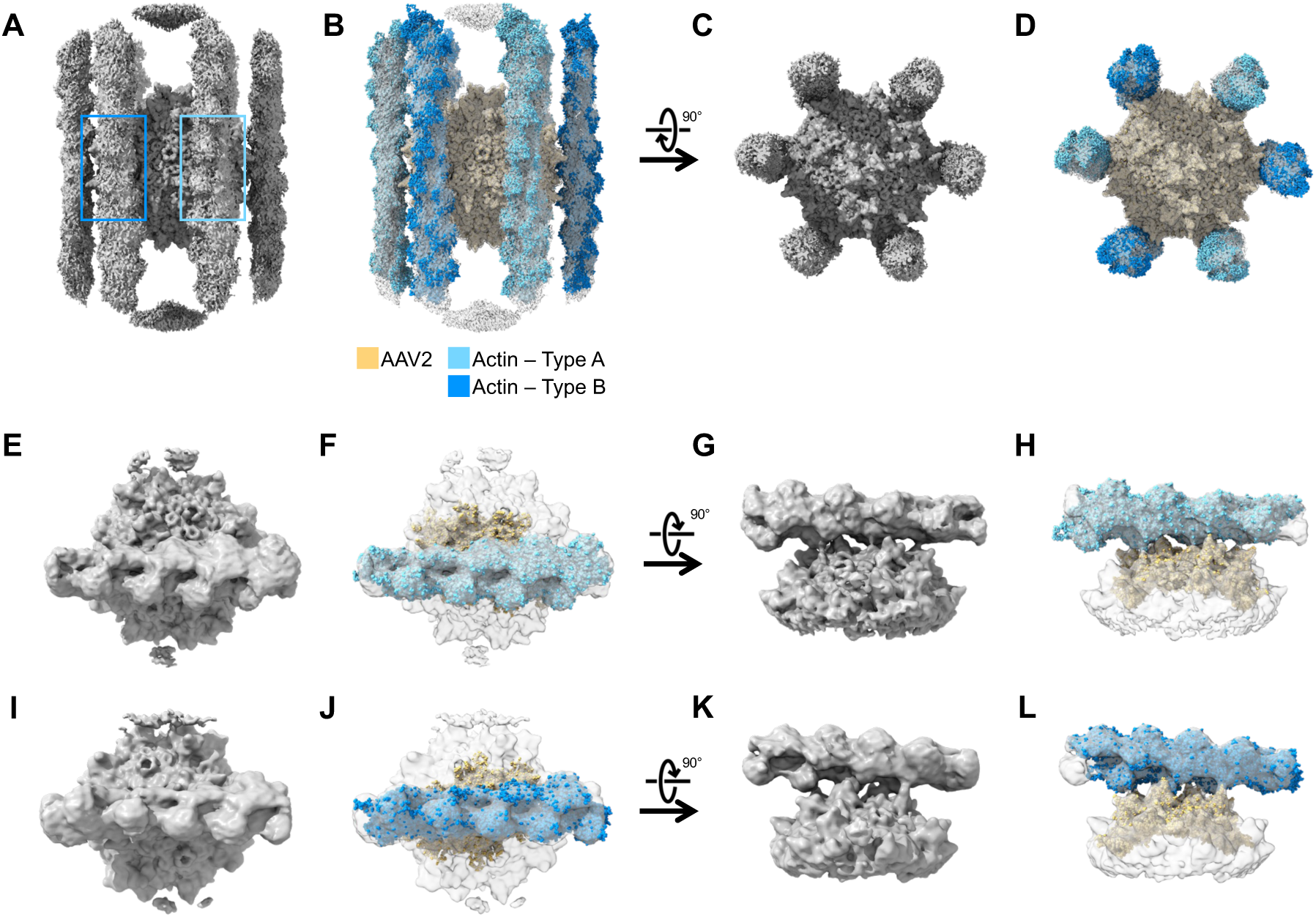
Reconstructions of AAV2 interacting with actin filaments. (A) 5.0 Å C3 reconstruction of actin filaments interacting with AAV2, contoured at 1.5 σ. The next and previous AAV2 capsid present in the bundle is seen at the upper and lower extremities. The light blue box denotes the region focused on for local reconstruction of the Type A interface and the dark blue box denotes the region focused on for local reconstruction of the Type B interface. (B) 60-mer model of AAV2 (orange) and six 16-mer models of actin filaments (light blue – Type A, dark blue – Type B) docked into the near-atomic reconstruction (transparent grey). (C) Same as in A, rotated 90° around the x axis. (D) Same as in B, rotated 90° around the x axis. (E) 5.2 Å C1 local reconstruction of the Type A AAV2-actin interface, contoured at 1.5 σ. (F) 6-mer model of AAV2 (orange) and 9-mer model of actin filament (light blue) are docked into the near-atomic reconstruction (light grey). (G) Same as in E, rotated 90° around the x axis. (H) Same as in F, rotated 90° around the x axis. (I) 5.3 Å C1 local reconstruction of the Type B AAV2-actin interface, contoured at 1.5 σ. (J) 6-mer model of AAV2 (orange) and 9-mer model of actin filament (dark blue) are docked into the near-atomic reconstruction (light grey). (K) Same as in I, rotated 90° around the x axis. (L) Same as in J, rotated 90° around the x axis.

To obtain accurate atomic models to dock into the reconstructed density for AAV2-actin complex, we determined the structure of AAV2 alone and actin filaments alone to 2.07 Å and 3.26 Å resolution, respectively (Supplementary Figure 5, Supplementary Figure 6A-B, and Supplementary Figure 7-8). Importantly, both structures overlayed with previously determined structures with RMSDs of less than 0.5 Å, indicating that there are no major structural rearrangements in KMEI buffer (Supplementary Figure 7G and Supplementary Figure 8E). Notably, our AAV2 structure is currently the highest resolution wild-type structure available, with visible electronic overlap between hydrogen bonds, and 28,080 spherical densities not attributable to the protein model, which were interpreted as water molecules (Supplementary Figure 8F-H).

### Atomic modeling of the AAV2-actin interfaces

We docked the atomic models into the full particle density map, using one copy of our AAV2 model and six copies of a 16-mer of our actin model (Figure 2B and D). All models fit the density well. The helical axis of two actin filaments directly opposite each other are approximately 330 Å apart, which corresponds to the AAV2 capsid diameter of 250 Å and the radius of two actin filaments (70 Å) and is consistent with our negative stain analysis. However, our full particle reconstruction did not account for possible differences in actin filament polarity. Therefore, to resolve the interface between AAV2 and actin, we used localized reconstruction (see Methods). We extracted sub-particles for Type A and Type B interfaces and performed 3D classification without alignment. This revealed that both Type A and Type B interfaces had actin filaments with opposite polarities. Type A interface had 42% (n=20,347) and 28% (n=13,735) well-defined sub-particles with the two opposing polarities (the remaining 30% of the sub-particles showed no clear actin density). Type B interface had 19% (n=9220) and 19% (n=9465) well-defined sub-particles with opposing polarities (the remaining 62% of the sub-particles showed no clear actin density). To average filaments with opposing polarities at both types of interfaces separately, we applied C2 symmetry relaxation during a second round of 3D classification. After local refinement, we obtained a reconstruction of the Type A and Type B interfaces at 5.2 Å and 5.3 Å nominal resolution, respectively (Figure 2E-L). We docked a 6-mer of AAV2 capsid proteins, which corresponds to two adjacent 3-fold protrusions, and a 9-mer of actin into each map. All models fit with high confidence, with a correlation coefficient of 0.92 for both types of actin–capsid interfaces and 0.88 for both AAV2 models. Interestingly, for both interfaces, the actin filament interdigitates tightly into the pair of 3-fold protrusions (Figure 2 E-L). The four protrusions closest to the 2-fold axis extend to either side of the actin filament and contact the surface of the filament. In both interfaces, the actin filament is oriented so the seam between the double helix is pointed towards the capsid surface.

We next determined potential contact residues on the surface of AAV2 or actin by selecting any residue within 5 Å of the other docked protein model, which accounts for a long hydrogen bond and 2 Å of rotamer rotation (Supplementary Figure 9 and Supplementary Table 2). Due to the resolution of the reconstruction, we cannot deterministically model side chain or loop shifts; therefore, the residues mentioned here are a conservative estimate, more residues may mediate the interaction. The actin filament contacts six monomers of AAV2 capsid protein in the Type A binding mode, and four monomers in the Type B binding mode. Five monomers of actin contact AAV2 in the Type A binding mode and four monomers contact AAV2 in the Type B binding mode. Interestingly, both conformations of actin interaction make contacts with AAV2 VR-I, -IV, -V, -VI, -VIII, and -IX. Several residues are shared between the interfaces that contact actin: T454, N587, Q589, and K706 (Supplementary Table 2, denoted by *). Additionally, the majority of residues in one binding interface are within three amino acids of residues within the other binding interface (Supplementary Table 2, denoted by §). However, some residues are unique to only one type of actin interaction (Supplementary Table 2). Likewise, some contact residues are shared between binding modes on the surface of actin, namely E100, S234, and S235 (Supplementary Table 3, denoted by *). In addition to these, many patches of residues are within three amino acids of residues on the other binding mode (Supplementary Table 3, denoted by §). While other interactions, are unique between Type A and Type B binding.

When we overlayed the Type A interface with the Type B interface, differences between the conformation of the actin filament were immediately apparent (Figure 3A-C). The two filaments are offset by 44° in the *xy* plane and the Type A filament is 5 Å closer to the surface of the capsid. Although the residue number of many contacts shared by Type A and Type B filaments are the same, some residues are in identical locations on the surface of the capsid, while others are on a different symmetry-related monomer (Figure 3D). For example, even though both types of filaments contact T454, the symmetry related monomer that is contacted differs. It is remarkable that the icosahedral symmetry of the AAV2 surface allows for two distinct actin binding conformations using similar residues. Likewise, while some contact residues on actin are shared, or nearby, the interaction interface on the surface of the actin filament is rotated approximately 180° between Type A and Type B (Figure 3E). Again, the helical nature of actin allows for unique interfaces using a similar set of residues.

**Figure 3.**
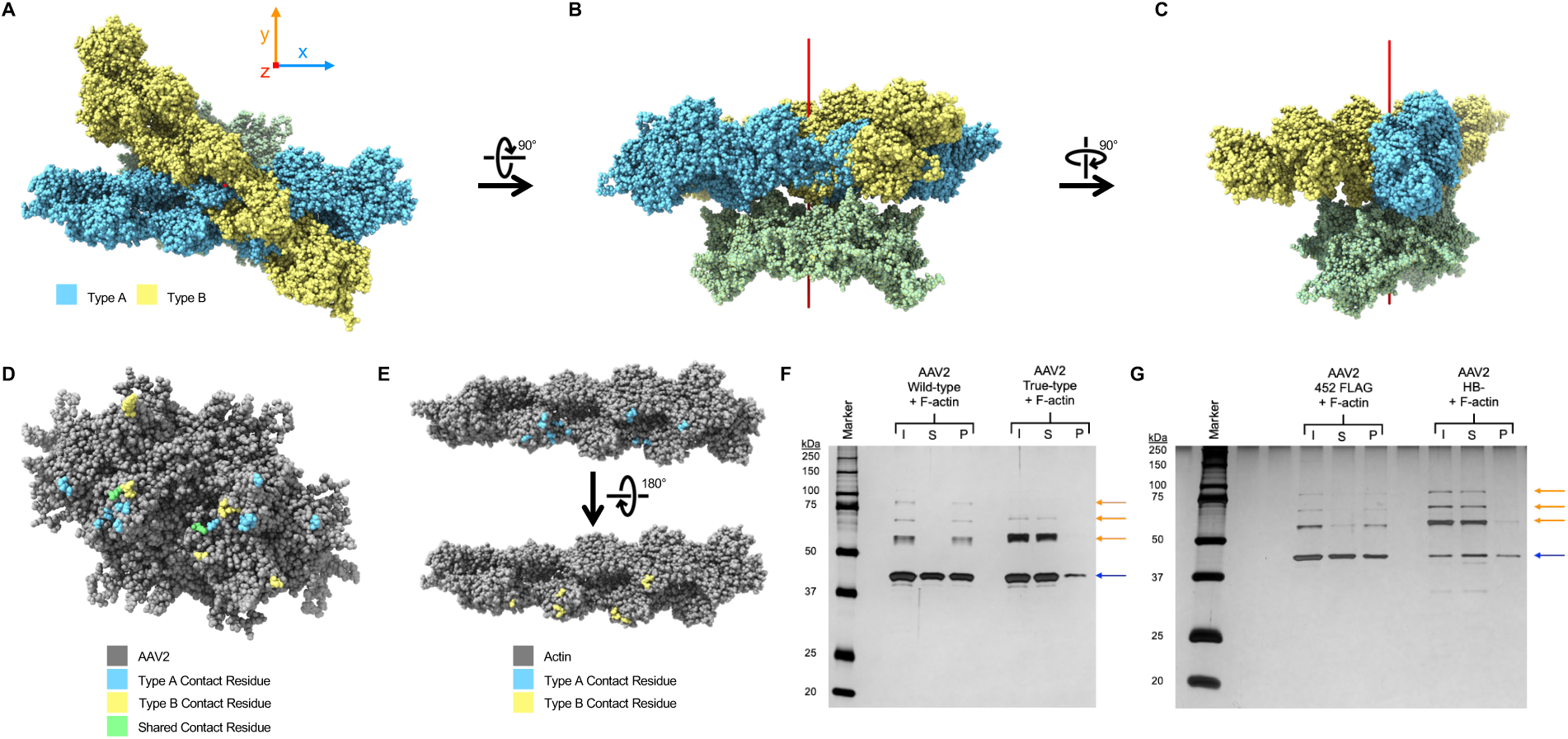
Actin filaments bind AAV2 in two poses but interactions are mediated by similar residues. (A) Overlay of the Type A AAV2-actin interface (light blue) and the Type B AAV2-actin interface (yellow). The 2-fold symmetry axis of AAV2 is oriented to be equivalent to the z axis and shown as a red rod. (B) Same as in A, rotated 90° around the x axis. (C) Same as in B, rotated 90° around the y axis. (D) 6-mer of AAV2 viewed down the 2-fold symmetry axis. Type A contact residues are colored light blue, Type B contact residues are colored yellow, and residues that are shared between Type A and Type B interfaces are colored green. (E) 9-mer of actin viewed perpendicular to the helical axis in two views, rotated 180° around the helical axis with respect to each other. Type A contact residues are colored in light blue and Type B contact residues are colored in yellow. (F-G) Silver stained SDS-PAGE gel containing fractions from a centrifugation-based actin bundling assay. “I” represents the input fraction, “S” represents the supernatant fraction, “P” represents the pellet fraction. The left group contains samples from the AAV2 wild-type + actin filament test reaction and the right group contains samples from the AAV2 true-type + actin filaments test reaction. AAV2 bands are indicated by orange arrows and actin bands are indicated by the blue arrow.

### AAV2 variants no longer bundles actin filaments

To validate the interaction site identified by our atomic docking studies, we repeated the centrifugation-based bundling assay using AAV2 true-type A593S (abbreviated as AAV2 TT hereafter), a previously characterized AAV2 variant that combines 13 of the most common amino acid differences found in circulating human population AAV2s into a single capsid, that differ from the clinically used AAV2 capsid.^42^ Importantly, AAV2 TT contains two variants that contact both the Type A and Type B actin filaments in our atomic models, R585S and R588T (Supplementary Table 2). Furthermore, AAV2 TT contains four mutations that are near to putative contact residues: Q457M, S492A, E499D, and F533Y (Supplementary Table 2). Interestingly, AAV2 TT does not bundle actin filaments, providing support for the interaction interfaces determined by our atomic docking studies (Figure 3F). It should be noted that AAV2 TT also contains seven other residues that are not located at the interface. However, five of these residues are on the interior of the capsid, so they are not expected to influence actin binding. G546D and E548G are on the exterior of the capsid, but they are not near the actin filament in either binding interface determined by our atomic docking.

To narrow down the key residues that mediate the AAV2-actin interaction, we repeated the bundling assay with two more targeted AAV2 variants. In the first, a FLAG peptide sequence (DYKDDDDK) was inserted in between residues 452 and 453, lengthening the 3-fold protrusions. We refer to this variant as AAV2 452-FLAG. In the second, we mutated R585S and R588T, which has been previously shown to disrupt heparin binding.^42^ We refer to this variant as AAV2 HB-. Interestingly, AAV2 452-FLAG is split between the supernatant and pellet fraction, indicating that it can still bundle actin filaments, but not as strongly as wild-type (Figure 3G). However, AAV2 HB- is found almost entirely in the supernatant, indicating that mutation of just these two key residues nearly ablates actin filament bundling (Figure 3G).

### The actin contact residues on the AAV2 capsid overlap with other receptor binding sites

The 3-fold protrusions also mediate interactions with two other important receptors/attachment factors: the site for heparan sulfate proteoglycan (HSPG) binding and the site for adeno-associated virus receptor (AAVR) binding.^43–48^ HSPG is expressed on the surface of most cell types, and it is thought to facilitate initial capsid attachment to the cell surface cell.^43–46^ AAVR is a membrane protein which recognizes many AAV serotypes, followed by shuttling to the trans-Golgi network.^47,48^ Both HSPG and AAVR are required for efficient cell transduction by wild-type AAV2. Interestingly, three residues that contact HSPG, K532, R585, and R588 also contact actin filaments (Figure 4A-B and Supplementary Table 2). Moreover, residues near the HSPG binding pocket, especially N587 and Q589, make multiple contacts with actin filaments in both binding modes. Importantly, R585 and R588 are shared between HSPG and actin filament binding, but differ in AAV2 TT and AAV2 HB-, supporting the dual binding capacity of these two residues, as neither variant bundles actin filaments or binds HSPG (Figure 4A-C and Supplementary Table 2).^42^ Two residues contact both actin binding modes as well as AAVR, Q589 and K706 (Figure 4A-B and Supplementary Table 2). Additionally, five other residues contact both AAVR and one binding mode of actin, while nine other AAVR contact residues are within three amino acids of actin contact residues (Figure 4A-B and Supplementary Table 2).

**Figure 4.**
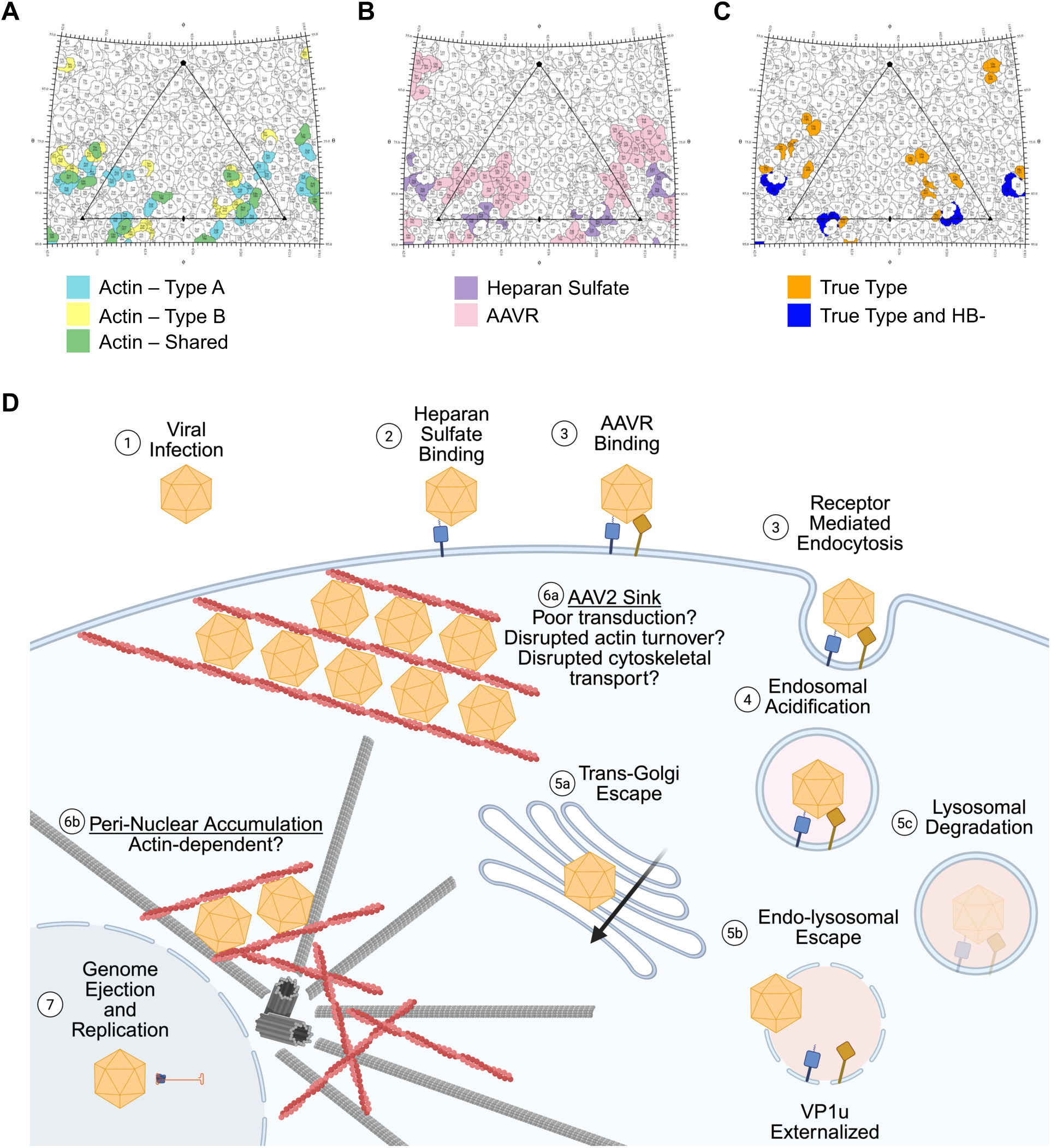
Impact of actin-binding on the transduction pathway of AAV2. (A) Stereographic roadmap projection of the surface of AAV2. The asymmetric unit is contained within the triangle, with icosahedral symmetry axes denoted by the black pentagon (5-fold), triangle (3-fold), and oval (2-fold). Type A actin contact residues are colored light blue, Type B actin contact residues are colored yellow, and contact residues shared between both interaction interfaces are colored green. (B) Same as in A, but Heparan Sulfate contact residues are colored purple and AAVR contact residues are colored pink. (C) Same as in A, but residues that are mutated in AAV2 true-type are colored orange and mutations that are shared between AAV2 true-type and AAV2 HB- are colored dark blue. (D) Cartoon depicting the transduction pathway of AAV2 into the nucleus of cells. Potential biological functions of the AAV2-actin interaction are shown in 6a and 6b.

We next compared the AAV2-interaction residues on actin filaments to residues that mediate interactions with other actin binding proteins (Supplementary Table 3). Interestingly, many actin contact residues also interact with multiple classes of actin binding protein, including other actin bundling proteins, actin severing proteins, actin polymerizing proteins, actin depolymerizing proteins, actin branching proteins, cytoskeletal adaptor proteins, and myosin motor proteins (Supplementary Table 3, shared residues denoted with ‡, nearby residues denoted by ◊). The wide range of proteins which share contact residues with AAV2 suggests that binding by AAV2 could have a significant impact on global cytoskeletal dynamics and function.

### Cryo-electron tomograms reveal a hexagonal lattice of AAV2 capsids among heterogeneous actin filaments

To visualize AAV2-actin bundles in 3D space at nanometre resolution, we utilized *in vitro* cryogenic electron tomography (cryo-ET). Tomograms were subjected to neural network-based segmentation, resulting in clearly distinguishable AAV2 capsids and actin filaments (Supplementary Figure 10 and Supplementary Video 3-4). We observed a wide range of AAV2-actin bundles, including very dense tangles where the filaments are overlapping and running in multiple directions, as well as some AAV2-actin bundles that were extremely ordered, with co-linear actin filaments (Supplementary Figure 10B-C *versus* 10D). Interestingly, AAV2 capsids are observed to bind to a heterogenous number of actin filaments, binding at least one filament to as many as six. Capsids are also bound to both straight filaments and bent filaments, including those with a near 180° turn (Supplementary Figure 10B). AAV2-actin “lattices” are found throughout our tomograms, where multiple parallel AAV2-actin bundles meet and form a multi-layer bundle (Supplementary Figure 10D, red box). Here, individual capsids form an almost crystalline hexagonal array, with approximately 360 Å between capsids in each direction, including bundles on top or below the main array (Supplementary Figure 10E-H). Indeed, global analysis of ten tomograms reveals that the distance between most capsids and their nearest neighbour is 300-400 Å (Supplementary Figure 11I). It is striking how despite the heterogeneity in the number of bound actin filaments and the curvature of those filaments, the distance between capsids in an AAV2-actin bundle is uniform.

### AAV2 does not bind monomeric actin

Since each actin monomer individually makes a limited number of contacts with AAV2, we did not expect monomeric actin to bind AAV2. The strong binding observed for actin filaments is likely mediated by multivalent interaction summed over multiple actin monomers. Since monomeric actin is only 42 kDa, which is approximately 1% of the molecular weight of AAV2, we would not be able to reliably detect a change in molecular weight upon binding, so centrifugation-based assays could not be used. Besides, negative stain images would not have the resolution required to distinguish bound monomeric actin. Therefore, we tested the interaction by incubating AAV2 in the presence of 120 times molar excess of monomeric actin, then plunge freezing the sample and collecting cryo-EM data. In both 2D class averages and the final 2.32 Å reconstruction, monomeric actin is not observed bound to the surface of AAV2 (Supplementary Figure 11). We cannot completely rule out the lack of interaction, as we did not exhaustively test different buffer systems and concentration ranges. However, if the interaction exists, it is much lower affinity than the actin filament binding affinity.

## Discussion

Here, we demonstrate that AAV2 specifically binds and bundles actin filaments *in vitro*. To the best of our knowledge, this is the first time a viral capsid has been observed to directly bundle actin filaments. Other viruses have been observed modulating actin dynamics to induce bundling. But to date, these viruses all rely on modulating the host signaling pathway or through an accessory protein.^27–31^ AAV2 is distinct from other actin bundling proteins since it has icosahedral symmetry and is made up of 60 copies of the capsid protein. Most other actin bundling proteins are either monomeric with multiple actin binding domains, or form small oligomers such as dimers.^25,49^ Therefore, we propose that AAV2 is the founding member of a new class of actin binding proteins, the actin bundling viruses (ABV).

The interaction between AAV2 and actin filaments could have wide-reaching implications in the use of AAV2 as a gene therapy vector (Figure 4F). First, it is possible that the bundling of actin filaments by AAV2 results in errors in intracellular trafficking or cell morphology when cells are transduced by AAV2. It would be interesting to visualize actin *in cellulo* using fluorescent microscopy to determine if AAV2 particles bundle *in cellulo*, and if so, whether this exogenous bundling causes measurable outcomes on cytoskeletal trafficking. Next, it is possible that actin filaments serve as an unknown barrier to successful transduction by AAV2. Though not explicitly measured here, the interaction between AAV2 and actin filaments appears robust, with nearly all of AAV2 binding and bundling actin filaments in the bundling assay, even though the concentration of AAV2 was 200 times lower than the concentration of actin filaments (Figure 1B). While the majority of actin filaments are intracellular, some actin filaments are found extracellularly, and levels of extracellular actin are heightened during disease and necrotic cell death.^50^ Therefore, it is possible that extracellular actin filaments limit the transduction efficiency of AAV2, since actin binding would preclude binding to the cell surface attachment factor, HSPG, or the cellular receptor, AAVR. Additionally, since actin is highly expressed in all cell types, and is especially highly expressed in muscle fibers, it is possible that upon intravenous injection of therapeutic AAV2 capsids, many of the vectors remain trapped after endo-lysosomal escape and bound to an actin filament “sink”. If this is true, it could explain why extremely high MOIs are required for therapeutic efficacy and would suggest that modulation of this binding interface between AAV2 and actin filaments may increase transduction efficiency. It is also possible that the durable gene expression observed upon rAAV2 transduction could be partly due to slow release of capsids from AAV2-actin “sinks” over time. Actin filaments localize to just inside the plasma membrane in adhesion junctions (Figure 1A).^51,52^ Therefore, it is possible that AAV2 is trapped just inside the cell membrane in these fluorescent microscopy experiments. However, due to the resolution of light microscopy, it is also possible that AAV2 capsids remain outside the membrane or bound to the membrane. Therefore, in the future, higher resolution methods like *in situ* cryogenic electron tomography (cryo-ET) are needed. Finally, a recent study showed that cytosolic actin is commonly found as a contamination during rAAV2 purification, suggesting that the interaction occurs in cells, and is strong enough to last through purification.^53^

However, the alternate hypothesis is also possible; actin filaments may serve to help AAV2 traffic to the nucleus. It was observed previously that destabilization of microtubules before AAV2 infection decreases the transduction efficiency by approximately two-fold but does not stop transduction completely.^16^ Actin-mediated transport seems to be the obvious alternative choice for trafficking, and their interaction as demonstrated here supports this hypothesis. AAV2 still accumulates in the peri-nuclear region, even when microtubules are depolymerized.^16^ Therefore, peri-nuclear accumulation of AAVs may be due to interactions with actin in the MTOC, not due to interactions with microtubules, as previously thought. Further supporting the importance of actin filaments in AAV2 trafficking is that when microtubules are depolymerized after the majority of capsids have made it to the peri-nuclear region, the transduction efficiency increases in a manner dependent on the RhoA-ROCK-mediated actin filament rearrangement.^17^ In this study, when only actin filaments were depolymerized, no effect on transduction was observed. However, this was done after AAV2 capsids were already trafficked to the peri-nuclear region. Since AAV trafficking is a complex, multi-step pathway, it will be important to unweave the interplay between microtubule- and actin-based trafficking during each step of the viral transduction pathway. It will be interesting in the future to determine whether actin filaments or myosin motors are required for transduction by AAV2.

Our cryo-ET results demonstrate the heterogeneity of the AAV2-actin interaction, as each capsid can bind a varying number of filaments, each filament can adopt various curvature, and each filament has intrinsic polarity. Actin is often bound intracellularly by many other proteins, including other bundling proteins. Therefore, the flexibility of this interaction likely allows for the engagement to take place under the various conditions found within the cell. In the future it will be interesting to use *in situ* cryo-ET to visualize AAV2 binding to actin in the context of the cell, where AAV2 will likely have to compete for binding with the myriad other actin binding proteins. Additionally, it is possible that the binding between AAV2 and actin is differs based on sub-cellular localization, so cryo-ET studies will be aimed at the peri-nuclear region and the periphery of the cell, among others. Furthermore, since other actin bundling proteins result in unique spacings between co-linear actin filaments, it will be interesting to see which, if any, of these actin bundles are still competent to bind AAV2. On the other hand, certain myosin motors require the unique spacing of a particular bundling protein, so it will be interesting to see if the wide spacing caused by AAV2 bundling are still competent for cytoskeletal trafficking.^54,55^

The structure of AAV2 interacting with actin filaments offers many interesting insights. First, the presence of two binding modes indicates that the interaction between AAV2 and actin filaments demonstrates that AAV2 can bind either side of the actin helix, since Type B actin is rotated ∼180° from Type A actin. The contacts between multiple monomers of AAV2, located in two sequential 3-fold protrusions, and multiple monomers of actin likely indicate that the interaction is strong, being at least bipartite. Since AAV2 was observed to bind as many as six actin filaments, the combined interaction interface is likely larger, indicating a multiplicative effect on binding affinity. Since actin filaments are locally constrained to their persistence length, we expect that once one AAV2 capsid binds, the local filament spacing would be ideal for a second capsid to bind. When this is repeated, cooperative bundling results, in line with what we observed in time-lapse negative stain experiments (Supplementary Figure 2). In the future, it would be revelatory to measure the binding affinity of AAV2 for actin filaments, as well as to determine if the cooperativity we propose here can be measured.

Perhaps most exciting, the structure of AAV2-actin localizes the putative interaction site to residues in surface exposed variable regions (VRs). The amino acids in these regions vary extensively between \ AAV serotypes and are responsible for cell binding and tropism.^15^ Therefore, it is possible that other AAV serotypes also bind and bundle actin filaments, perhaps with even higher affinity, hence future capsid variant designs should study actin bundling phenotypes. Residues T454, found in VR-IV, as well as R585, N587, R588 and Q589, found in VR-VIII, were observed to mediate multiple contacts between AAV2 and actin filaments. This is particularly interesting because VR-IV and VR-VIII have been the sites of extensive capsid engineering efforts, since they can be extensively mutated and still assemble into functional capsids. ^1,3–5^ It will be interesting to re-evaluate the effect AAV2 variants have on transduction considering the actin interaction. Indeed, AAV2 TT, which harbors 13 amino acid substitutions, no longer bundles actin filaments, confirming the binding site functionally. Additionally, AAV2 HB- does not bundle actin, implicating just two key residues that mediate the interaction. It is interesting that AAV2 452-FLAG still partially bundles actin filaments. Both binding modes have a 3-fold protrusion interdigitated into the actin seam, so we thought that extending the 3-fold protrusions would prevent bundling entirely. Unfortunately, our two actin-binding deficient AAV2 variants, AAV2 TT and AAV2 HB-, harbor two mutations in R585 and R588 that are known to be important for heparan sulfate binding, which mediates cell surface attachment.^42^ Therefore, we cannot deconvolve the impact of heparan sulfate binding with the impact of actin binding on transduction efficiency using these capsid variants.^42^ Therefore, we suggest future research efforts in capsid design should be devoted to identifying specific mutations that disrupt the AAV2-actin interaction but do not disrupt the heparan sulfate interaction to separate the functions of each interaction on the transduction efficiency of AAV2. Perhaps by inserting a longer, more rigid, or less charged peptide at residue 452 to block the AAV2-actin interaction. It will be noteworthy to see what effect on transduction efficiency an actin-binding deficient mutant will have. Transduction efficiency increasing would support the hypothesis that actin filament binding acts as a “sink”, soaking up capsids on their way to the nucleus. Transduction efficiency decreasing would support the hypothesis that AAV2 traffics on actin filaments on the way to the nucleus or accumulates in the peri-nuclear region due to actin binding. Either way, the biological role of the AAV2-actin interaction is sure to be critical, and modulating the AAV2-actin interaction could lead to the development of a better gene therapy vector.

Finally, the robust interaction between AAV2 and actin filaments has interesting implications for the biotechnology sector. Namely, the interaction can be repurposed as a purification method.^53^ After AAV2 expression in packaging cells, the cell lysate is clarified and much of the AAV2 is in the pellet, not the supernatant as would be expected based on its molecular weight. However, washing the pellet with 500-1000 mM sodium chloride liberates the AAV2 from the pellet. Considering the AAV2-actin interaction, this purification phenotype is consistent with AAV2 interacting with endogenous actin and the high sodium chloride being required to break up the AAV2-actin bundles. However, actin can be readily polymerized and depolymerized through buffer exchanges or through the addition of actin-depolymerizing or filament-stabilizing small molecules. Therefore, after liberation of rAAV2 from the pellet, the supernatant separated and exogenously purified actin can be added in. Actin can then be polymerized and the resultant AAV2-actin bundles can be isolated by a low-speed centrifugation step. Finally, the actin filaments can be resuspended and depolymerized then rAAV2 can be separated from monomeric actin by a simple filtration step, since AAV2 does not bind monomeric actin (Supplementary Figure 11). This method of purification should be rapidly scalable and economical, since large quantities of actin can be purified from rabbit or chicken muscle acetone powder at low cost. Furthermore, with fine tuning, it should be possible to leverage endogenous actin for purification of AAV2 using buffer exchanges and low-speed centrifugation alone.

Taken together, the discovery that AAV2 binds and bundles actin filaments has many paradigm-altering implications both in basic AAV biology and in applied AAV-mediated gene therapy. Future study of this interaction will be critical to understand how to leverage it for next-generation AAV design with improved transduction efficiency and lower off-target effects, better serving the patients that rely on these life-saving treatments.

## Methods

### Production of adeno-associated virus 2

To generate adeno-associated virus 2 (AAV2) virus-like particles (VLPs), HEK293 cells were transiently transfected with two plasmids, pXR2 and pHelper. pXR2 expresses the AAV2 *Rep* and *Cap* open reading frames (ORFs) and pHelper expresses four genes from adenovirus which are necessary for AAV production.

HEK293 cells were seeded in 15 cm plasma-treated tissue culture dishes in 20 mL of DMEM + 10% FBS. When the cells reached ∼90-100% confluency, the media was exchanged for 15 mL of fresh, serum-free DMEM and cells were allowed to recover for 1 hour in an incubator at 37°C, 5% CO_2_. Transfection reagent was prepared by adding 13 μg pXR2, 20.5 μg pHelper, and 125 μL polyethylenimine (pre-warmed to 56°C) to 1 mL Opti-MEM (pre-warmed to 37°C) for each 15 cm plate to be transfected. Transfection reagent was often prepared in 10 plate batches. After 15 minutes of incubation at room temperature, the transfection reagent became cloudy, due to the formation of polyethylenimine-plasmid DNA assemblies, and 1120 μL of transfection reagent was added to each 15 cm plate. Cells were incubated at 37°C, 5% CO_2_ for 3-4 hours to allow uptake of the plasmids. Finally, 5 mL of DMEM + 10% FBS was added to each plate, and cells were incubated at 37°C, 5% CO_2_ for 72 hours to produce AAV2 VLPs.

After 72 hours, cells were harvested by using a sterile cell scraper and transferred into sterile 500 mL centrifuge bottles. Cells were pelleted by centrifugation at 1500 rpm in a Beckman Coulter JA-10 rotor for 20 minutes at 4°C. The supernatant was poured off into sterile 1000 mL glass media bottles for later processing. Cells were resuspended in 1 mL/plate of 1X TD (1X PBS, 5 mM MgCl_2_, 2.5 mM KCl, pH 7.4), vigorously vortexed, and flash frozen in liquid nitrogen. Frozen cell pellets were kept at -20°C until ready for purification.

10% (w/v) PEG8000 was added to the supernatant and was incubated at 4°C with constant stirring for 72 hours. Then, PEG-AAV assemblies were pelleted by centrifugation at 9000 rpm in a Beckman Coulter JA-10 rotor for 90 minutes at 4°C. The supernatant was discarded and the pellet was resuspended in 1 mL/10 plates of 1X TD (1X PBS, 5 mM MgCl_2_, 2.5 mM KCl, pH 7.4). Resuspended PEG-AAV2 was frozen at -20°C until ready for purification.

### Purification of adeno-associated virus 2

When ready for purification, cells were lysed by three freeze-thaw cycles. After the final thaw, 25 units of benzonase were added for every 10 plates of cells which went into the purification. Cell lysate was incubated with benzonase for 1 hour at 37°C to digest nucleic acids. The cell lysate was clarified by centrifugation at 10,000 rpm in a Beckman Coulter JA-20 rotor for 30 minutes at 4°C. The cell lysate supernatant was transferred to a sterile 50 mL conical tube. The pellet was washed with 5 mL of 0.8X TD + 1 M NaCl, made by the addition of 20% (v/v) 5 M NaCl to 1X TD, and clarified again by centrifugation at 10,000 rpm in a Beckman Coulter JA-20 rotor for 30 minutes at 4°C. The salt wash supernatant was transferred to a sterile 50 mL conical tube and diluted up to 50 mL with 1X TD to bring the concentration of NaCl to ∼200 mM. 1 mL of AVB Sepharose High Performance affinity resin (pre-equilibrated in 1X TD) was added to both the cell lysate supernatant and the diluted salt wash supernatant and incubated overnight at 4°C with constant mixing using a conical tube rotator.

The following morning, the AVB resin was pelleted by centrifugation at 1,000 x *g* for 1 minute at 4°C. The supernatant was poured off and the AVB resin was resuspended in 10 mL of 1X TD to remove any non-specifically bound proteins. The AVB resin was then pelleted again and the wash step was repeated. To elute AAV2 VLPs the pelleted AVB resin was resuspended with 9 mL of AVB Elution Buffer (0.1 M glycine, pH 2.7) and incubated for 5 minutes. The AVB resin was then pelleted and the supernatant was carefully transferred using a serological pipette to a conical tube containing 0.9 mL of AVB Neutralization Buffer (1 M Tris Base, pH 10). The tube was gently inverted to neutralize the highly acidic elution buffer. AAV2 VLPs were dialyzed against 4 L of 1X PBS pH 7.4 for at least 6 hours. Dialysis was repeated for a total of 3 times.

To concentrate AAV2 VLPs, the dialyzed solution was evenly divided into two 5 mL ultracentrifuge tubes and balanced to ± 0.01 g. AAV2 VLPs were pelleted by centrifugation at 50,000 rpm in a Beckman Coulter SW55Ti rotor for 75 minutes at 4°C. The supernatant was discarded and pellets were resuspended in ∼100-200 μL of KMEI (50 mM KCl, 1 mM MgCl_2_, 1 mM EGTA, 10 mM imidazole pH 7.0). Resuspended virus was further dialyzed against 200 mL of KMEI three times to ensure the final virus preparation was in KMEI. Concentrated, purified virus was stored in aliquots at - 80°C. Thawed aliquots were used within four weeks after thawing.

The purity and concentration of AAV2 VLPs was determined by running 5 μL of virus on a 10% SDS-PAGE gel alongside BSA standards. The gel was stained with GelCode Blue overnight and destained for several hours in ddH_2_O. The gel was then imaged using a BioRad ChemiDoc on colorimetric mode. Purity was assessed by the absence of contaminating protein bands. A standard curve was generated by densitometric quantification of the BSA standards and was used to determine the concentration of AAV2 VLPs.

### Actin purification and preparation

Rabbit skeletal α-actin was purified from rabbit muscle acetone powder as previously described.^56^ Briefly, 5 grams of rabbit muscle acetone powder (Pelfreez) was rehydrated in 100 mL of G-buffer (2 mM Tris HCl pH 8.0, 0.2 mM Na_2_ATP, 0.1 mM CaCl_2_, 1 mM NaN_3_, 0.5 mM DTT) and stirred at 4°C until gelatinous. Unextracted tissue was pelleted by centrifugation at 32,000 x *g* in a Beckman Coulter Ti70 rotor for 35 minutes at 4°C. The supernatant was poured through a cheese cloth to remove small pieces of tissue which did not pellet. The volume of the solution was measured in a graduated cylinder and then adjusted to 50 mM KCl and 2 mM MgCl_2_ with stock solutions. Polymerization of extracted globular actin (G-actin) was allowed to proceed with gentle stirring for 60 minutes at room temperature. After 60 minutes, the solution was viscous and trapped air bubbles, indicating that the solution was polymerized to actin filaments. To dissociate tropomyosin, solid KCl was added to the solution slowly until the final concentration of KCl reached 800 mM (50 mM in polymerization, 750 mM added assuming no volume change). The solution was then stirred at room temperature for a further 30 minutes followed by 30 minutes of gentle stirring at 4°C. Actin filaments were then pelleted by centrifugation at 141,000 x *g* in a Beckman Coulter Ti70 rotor for 150 minutes at 4°C. The supernatant was discarded and the gelatinous pellet was washed gently with G-buffer twice. The pellet was then scrapped into a Dounce homogenizer, 2 mL of G-buffer was added, and then the pellet was homogenized by several depressions of the loose plunger. Resuspended actin filaments were added to a 12-14 kDa MWCO dialysis membrane and dialyzed against 2 L of G-buffer three times. After dialysis, monomeric G-actin was separated from filaments that did not fully depolymerize by centrifugation at 149,000 x *g* in a Beckman Coulter SW55Ti rotor for 150 minutes at 4°C. The top two thirds of the supernatant is monomeric G-actin and was aspirated, aliquoted, flash frozen in liquid nitrogen, and stored at -80°C for until needed for further use. The bottom third was discarded.

To make actin filaments for use in bundling assays and electron microscopy, an aliquot of G-actin was thawed at 4°C, then KCl stock was added to a final concentration of 50 mM and MgCl_2_ stock was added to a final concentration of 1 mM. The solution was then placed in 12-14 kDa MWCO dialysis membrane and dialyzed against 1 L of Polymerization Buffer (10 mM MOPS pH 7.0, 50 mM KCl, 2 mM MgCl_2_, 0.5 mM EGTA, 0.2 mM ATP, 1 mM DTT, 1 mM NaN_3_) overnight at 4°C. The next morning, the dialysis membrane was transferred to 1 L of Actin Storage Buffer (4 mM MOPS pH 7.0, 2 mM MgCl_2_, 0.1 mM EGTA, 1 mM DTT, 3 mM NaN_3_) and dialyzed for 8 hours. This dialysis was repeated twice, for a total of three sequential dialyses against Actin Storage Buffer. Finally, the concentration of actin filaments was measured using its UV absorbance at 290 nm using an extinction coefficient of 26,600 M^-1^cm^-1^. Actin filament preps were typically between 120-160 μM. The purity of the actin filament preps was confirmed by running 2 μg on an SDS-PAGE gel. Actin filaments were stored at 4°C for up to a month before use.

For TIRF polymerization experiments, rhodamine-labeled actin (AR05) was obtained from Cytoskeleton, Inc. and hydrated according to the manufacturer’s instructions. Following hydration, actin was dialyzed overnight against 2 L of G-buffer (concentrations in mM unless otherwise noted): Tris-HCl (2, pH 8.0), ATP (0.2), CaCl₂ (0.1), DTT (1), NaN₃ (1). To remove actin aggregates and preformed nuclei, dialyzed actin was clarified by ultracentrifugation at 100,000 × g for 60 min at 4 °C. The supernatant was recovered and used for polymerization assays.

### Immunofluorescence confocal microscopy

HeLa cells (ATCC® CCL-2™) were subcultured in 35 mm glass-bottom dishes (MatTek Corp.) at a density of 1 × 10⁶ cells per well. Cell confluency was monitored prior to infection. Cells were infected either without virus (control), with AAV2-GFP, or with AAV2 virus-like particles (VLPs). Following infection, cells were incubated for 30 minutes at 4°C. After incubation, cells were immediately washed with cold PBS, and this time point was designated as “0 min.” Cells were then further incubated at 37°C for up to 2 hours.

Following treatment and infection, media was removed, and cells were washed with 1 mL cold PBS. Cells were fixed with 4% paraformaldehyde for 30 minutes at 4°C, followed by three washes with cold PBS. Cells were then permeabilized with 0.2% Triton X-100 in PBS for 15 minutes at 4°C and washed three times with PBS.

Non-specific binding sites were blocked by incubating cells with 5% normal goat serum (NGS) in PBS for 1 hour at room temperature. After washing with PBS, cells were incubated overnight at 4°C with primary antibody (mouse monoclonal anti-A20; 1:500 dilution). Following overnight incubation, cells were washed three times with PBS and incubated with secondary antibody (goat anti-mouse Alexa Fluor 546) for 2 hours at room temperature.

Cells were then washed with PBS and counterstained for the actin cytoskeleton and nucleus using ActinGreen™ 488 (1 drop/mL) and NucBlue™ (1 drop/mL), respectively. After staining, cells were washed with PBS, mounted using ProLong Glass Antifade Mountant, and covered with a round glass coverslip.

Fluorescently labeled cells were imaged and analyzed using a confocal laser scanning microscope (Zeiss LSM 800; Zeiss, Jena, Germany). Imaging was performed using laser wavelengths of 405 nm for NucBlue™ (DAPI), 488 nm for ActinGreen™, and 543 nm for A20 labeled with Alexa Fluor® 546. Images were acquired using a Plan-Apochromat 63×/1.40 oil DIC M27 objective under identical laser exposure and microscope settings across all samples, using ZEN 2010 software. Post-acquisition image processing involved uniform adjustment of brightness and contrast across all images within each experimental set.

### Actin bundling assay

To measure the ability of AAV2 to bundle actin filaments, a centrifugation-based bundling assay was used. First, actin filament stock was diluted to 10 μM in KMEI (50 mM KCl, 1 mM MgCl_2_, 1 mM EGTA, 10 mM imidazole pH 7.0) and centrifuged at 15,000 x *g* for 1 hour at 4°C to pellet any spontaneously formed actin bundles. Following the spin, the top two thirds was transferred to a new tube and the concentration was measured using the absorbance at 290 nm (ε_290_ = 26,600 M^-1^ cm^-1^). Single actin filaments were then diluted to 2 μM in KMEI. 15 μL of 2 μM single actin filaments were then mixed with 15 μL of the following in separate reaction tubes: KMEI, 2 μM BSA in KMEI, KMEI adjusted to 30 mM MgCl_2_, and 10 nM AAV2 VLPs in KMEI. Reaction mixtures were mixed gently by pipetting then incubated at 4°C for one hour. After incubation, 15 μL was transferred to a clean microcentrifuge tube as the input fraction. The remaining 15 μL was centrifuged at 10,000 x *g* for 10 minutes at 4°C to pellet any actin bundles that formed during the reaction period. 15 μL of supernatant was carefully aspirated by using a pipette and transferred to a clean microcentrifuge tube. The pellet was resuspended with 15 uL of KMEI by gentle pipetting. 3 μL of 6X Laemmli gel loading dye (375 mM Tris HCl pH 6.8, 9% (w/v) SDS, 50% (v/v) glycerol, 9% (v/v) β-mercaptoethanol, 0.03% (v/v) bromophenol blue) was added to each 15 μL fraction and vortexed. Proteins were denatured by heating at 100°C for 5 minutes and all the solution was collected at the bottom of the tube by a brief spin in a microcentrifuge. 5 μL of denatured protein was loaded on a 10% SDS-PAGE gel which was run at 170 V for 70 minutes. After running, the gel was fixed and silver stained according to the manufacturer’s instructions (Pierce Silver Stain Kit). The stained gel was imaged using a BioRad ChemiDoc on white light transillumination mode.

### TIRF-based actin polymerization assay

High tolerance glass coverslips (24 × 50 mm, #1.5; Marienfeld Superior, Germany) were cleaned by sequential sonication for 10 min in 1-2% Hellmanex, rinsed three times with Milli-Q water, sonicated for 10 min in 100% ethanol, rinsed three times with Milli-Q water, and sonicated for an additional 10 min in Milli-Q water. Coverslips were dried under a nitrogen stream and plasma cleaned in argon for 5 min using a ZEPTO plasma cleaner (Diener Electronic, Germany). Plasma-cleaned coverslips were passivated by applying 100 µL Sigmacote (Sigma-Aldrich) to each coverslip and gently tilting to ensure complete surface coverage. Excess Sigmacote was immediately wicked away using a Kimwipe, and the coverslips were allowed to dry completely. After drying, coverslips were rinsed five times with Milli-Q water and dried under a nitrogen stream. Flow chambers were assembled by sandwiching double-sided adhesive tape between a microscope slide and a passivated coverslip to form a channel. Chambers were incubated with one chamber volume of 0.5% (w/v) Pluronic F-127 diluted in KMEID buffer (concentrations in mM unless otherwise noted): KCl (50), MgCl₂ (1), EGTA (1), imidazole (10), pH 7.0, DTT (1) for 15 min at room temperature. Excess Pluronic F-127 was removed by washing with two chamber volumes of KMEID buffer. Chambers were used immediately for experiments.

Actin polymerization assays were performed in Pluronic F-127 functionalized flow chambers using a final actin concentration of 1 µM, consisting of 20% rhodamine-labeled actin and 80% unlabeled actin. Polymerization was carried out in the presence or absence of 5 nM virus at 25 °C. Reactions were performed in polymerization buffer (concentrations in mM unless otherwise noted): Tris-HCl (2, pH 8.0), CaCl₂ (0.1), NaN₃ (1), ATP (0.2), DTT (0.5), KCl (50), MgCl₂ (2), EGTA (1), glucose (15), supplemented with catalase (20 µg·mL⁻¹), glucose oxidase (100 µg·mL⁻¹), and methylcellulose (0.2% w/v). Time-lapse imaging was performed using objective-style total internal reflection fluorescence microscopy and an inverted Nikon (Ti2-E) with motorized illuminator (H-TIRF) and a 100x oil objective (Nikon CFI Apochromat, N.A. 1.49). Samples were illuminated using a 561 nm laser line and emitted light (ET 630/75m, Chroma) captured on an EM-CCD camera (iXon Ultra 888, Oxford Instruments) controlled by NIS-Elements (AR v5.2, Nikon).

### Negative staining and imaging

Copper 400 mesh continuous carbon grids were glow discharged in air at 15 mA, 0.39 mbar for 25 seconds using a Pelco EasiGlow operating in negative mode. Hydrophilized grids were used within 30 minutes of glow discharge. 3 μL of sample (specified in each figure legend) was applied to each grid and incubated for one minute to allow particles to adsorb to the carbon surface. The grid was then washed to remove non-adsorbed particles by gently dabbing against a 15 μL drop of ddH_2_O five times. This wash step was repeated twice for a total of three wash drops. The water was then wicked away by tapping a torn piece of filter paper against the side of the grid. The grid was then stained by gently dabbing against 15 μL of freshly-filtered 1% (w/v) uranyl acetate five times. This staining step was repeated once for a total of two staining drops. Excess stain was wicked away by tapping a torn piece of filter paper against the side of the grid. The grid was then allowed to air dry until it was no longer visibly wet. Negatively stained grids were usually imaged the following day, but were stored in a grid storage box for up to six months.

### Negative stain data processing

Negative stain micrographs were converted to the MRC format using EMAN2 and then subjected to particle picking and 2D classification using CryoSPARC v4.7.1.^57,58^ Briefly, 756 particles were initially selected by manual picking and subjected to 2D classification. From this, five classes were used as templates to pick 31,996 particles using the template picker. After filtering out erroneous picks, 4877 particles were extracted using a box size of 256 pixels (2109 Å) and subjected to further 2D classification.

### Vitrification

To make AAV2-actin bundles for vitrification, AAV2 VLPs in KMEI were diluted to 60 nM and mixed in a 1:1 ratio with actin filaments diluted to 1 μM in KMEI, resulting in a final concentration of 30 nM AAV2 VLPs and 0.5 μM actin filaments. Additionally, to collect sparser micrographs, AAV2 VLPs in KMEI were diluted to 2 nM and mixed in a 1:1 ratio with actin filaments diluted to 400 nM, resulting in a final concentration of 1 nM AAV2 VLPs and 200 nM actin filaments. Complexes were allowed to form by incubation on ice for 90 minutes. Immediately prior to vitrification, complexes were gently mixed by pipetting. 3 μL of AAV2-actin bundles were pipetted onto a C-flat 1.2/1.3 holey carbon gold 300 mesh grid held within the sample chamber of a Leica EM GP2 plunge freezer, held at 95% humidity and 4°C. After 60 seconds of incubation, the grid was blotted from the back using Whatman no. 5 paper for 4 seconds and immediately plunged into liquid ethane.

To make actin filaments alone control grids, actin filaments was diluted to 1 μM in KMEI and vitrified as described in the preceding paragraph. Similarly, to make AAV2 VLP alone control grids, AAV2 VLPs in KMEI were diluted to 1 nM and vitrified as described in the preceding paragraph.

Grids were screened for ice thickness and particle distribution using a Jeol Cryo ARM 300 operating at 300 kV.

### Single particle data collection

Data was collected using a Jeol CryoARM300 (RRID:SCR_024555, SCR_020163) operating at 300 kV equipped with a Gatan K3 direct electron detector, a cold field emission gun electron source, and an in-column omega energy filter with a slit width of 20 eV. Exposures were recorded using SerialEM at a nominal magnification of 50,000X, resulting in a calibrated pixel size of 0.62 Å.^59^ Movies were fractionated into 50 frames using a total dose of 50 e^-^/Å^2^. Between 5 and 7 exposures were recorded per hole and 9 holes in a 3×3 array were imaged using image shift per stage translation. Defocus was targeted to be between 0.8 and 1.8 μm underfocus. 21,154 movies across two grids at multiple stage tilts were collected for AAV2-actin bundles, 1521 movies from one grid at 0° stage tilt were collected for AAV2 VLPs alone, and 1699 movies from one grid at 0° stage tilt were collected for actin filaments alone.

### Single particle data processing and 3D reconstruction

To reconstruct AAV2 VLPs alone in KMEI, CryoSPARC v4.7.1 was used.^57^ Briefly,1521 movies were motion-corrected and dose-weighted using the Patch Motion Correction module and initial CTF parameters were estimated using the Patch CTF Estimation module. Exposures with a maximum CTF fit greater than 10 Å were excluded from further processing, resulting in 1241 accepted exposures. Particles were initially picked using the Blob Picker module set to pick circular particles with a diameter of 230-270 Å, resulting in 105,258 particle picks. Particles were filtered using their local power score and normalized cross-correlation to filter out erroneously picked particles, resulting in 26,421 particles that were extracted in 500 pixel boxes and subjected to 2D classification. High-resolution class averages which resembled the expected icosahedral geometry were used as templates to pick 425,783 particles using the Template Picker module. Particles were filtered using their local power score and normalized cross-correlation to filter out erroneously picked particles, resulting in 37,526 high-confidence particles that were extracted using a 500 pixel box and subjected to 2D classification. High-resolution class averages which resembled the expected icosahedral geometry were used to create an Ab-Initio 3D reconstruction. The initial ab-initio model was low pass filtered to 30 Å and refined using Non-Uniform Refinement with icosahedral symmetry imposed.^60^ After refining per-particle defocus and per-group CTF parameters, as well as correcting higher order aberrations and for the curvature of the Ewald sphere, refinement converged to a 3D volume with a gold-standard Fourier shell correlation (GSFSC) masked resolution of 2.07 Å.^39^ The density map was sharpened using a negative B factor of -60.9, the slope of a linear fit to the Guinier plot. Reconstruction statistics are reported in Supplementary Table 1.

To reconstruct actin filaments in KMEI, CryoSPARC v4.7.1 was used.^57^ Briefly, 1699 movies were motion-corrected and dose-weighted using the Patch Motion Correction module and initial CTF parameters were estimated using the Patch CTF Estimation module. Exposures with a maximum CTF fit was greater than 10 Å were excluded from further processing, resulting in 1639 accepted exposures. 146 particles were manually picked and subjected to 2D classification to generate two initial low-resolution templates. These templates were used as input for the Filament Tracer module.^61^ The diameter of the filament was set to 70 Å and the separation between segments was set to 1.15 times the filament diameter (80.5 Å), which corresponds to three actin monomers. From these 81,548 segments were identified and filtered by local power score, curvature, and sinuosity, resulting in 56,733 segments chosen for extraction. Using a box size of 512 pixels, 52,984 segments were extracted and subjected to 2D classification. 48,980 segments contributed to featureful classes and were used in helical refinement with an initial helical rise of 28 Å, initial helical twist of -166°, an order of 3, and a dynamic mask clipped to 85% of the z axis.^62^ This produced an initial reconstruction at 3.64 Å. Local CTF parameters were refined and then refined particles were used in a second helical refinement using the same settings as previously. This resulted in a final reconstruction with a GSFSC masked resolution of 3.26 Å. The density map was sharpened using a negative B factor of -86.7. Reconstruction statistics are reported in Supplementary Table 1.

To reconstruct AAV2 VLPs interacting with actin filaments in KMEI, we performed a multi-step processing workflow detailed in Supplementary Figure 4, using CryoSPARC v4.7.1, Scipion3, and RELION5.^57,63,64^ Briefly, five datasets from two grids were collected at multiple stage tilts. For each dataset individually, micrographs were motion corrected and dose weighted using the Patch Motion Correction module, then CTF parameters were estimated using the Patch CTF Correction module in CryoSPARC v4.7.1. AAV2 capsids bound to actin filaments were manually selected then subjected to 2D classification. Classes that had the expected icosahedral geometry were selected and used as templates in template picking. Particles were filtered using their local power score and normalized cross correlation, then extracted using a box size of 1024 pixels and Fourier cropped to 256 pixels. After further 2D classification, particles were imported into Scipion3. After two rounds of 2D classification to filter out overlapping capsids, 155,468 particles were used to generate an icosahedral averaged reference volume. Using this volume, the capsid density was subtracted from the particles and subtracted particles were used in 2D classification. Classes that had clearly aligned actin filaments were selected (57,824 particles). These particles were subjected to three rounds of 3D classification with icosahedral symmetry relaxation in a custom version of RELION 5 supporting the relaxation of non-cyclic point group symmetries. In the first and second round, a 600-Å and 475-Å diameter spherical masks were used, respectively.. Actin filaments were observed to bind in a pseudo-6-fold manner around the icosahedral 3-fold axis. In the third round, a C6 rotational mask, generated from the actin filament densities, was used. The two classes with the best-resolved actin density were combined (16,313 particles) and the Euler angles of each subtracted particle was assigned to the respective unsubtracted particles. Particles were then rotated from the icosahedral to the C3 symmetry convention and subjected to 3D auto refinement with C3 symmetry enforced. Due to the inclusion of tilted datasets, the per-particle CTF was refined using CryoSPARC, resulting in a final C3 reconstruction at 5.0 Å, which is Nyquist limited. This reconstruction is referred to as the full particle reconstruction in the text.

In the full particle reconstruction, the polarity of actin filaments was not resolved, limiting the interpretability of the binding interface. Therefore, we reconstructed the interface between AAV2 and actin filaments using localized reconstruction. First, we picked subparticles for both Type A and Type B AAV2-actin binding modes and extracted them, resulting in 48,929 subparticles for Type A and 48,928 subparticles for Type B. These subparticles were used to reconstruct a reference volume for each type separately. Then, particles were subjected to 3D classification using a mask that included the entire actin density, and the very tips of the 3-fold protrusions. Three classes were selected for each Type A and Type B, resulting in 34,082 subparticles for Type A and 18,685 subparticles for Type B. Selected subparticles were then subjected to further 3D classification using C2 symmetry relaxation, again using the actin mask. From this, the 3D class with the most featureful density (Type A – 8,998 subparticles, Type B – 4,736 subparticles) was selected and used in CryoSPARC local refinement to reconstruct the entire interface with dynamic masking. Doing so resulted in a final reconstruction of the Type A interface at 5.2 Å and of the Type B interface at 5.3 Å. Final maps were sharpened by applying an inverse B-factor. Reconstruction statistics are shown in Supplementary Table 1.

### Model building and structure refinement

To build the atomic model of actin filaments, a high resolution cryo-EM structure (PDB: 8A2T) was rigid body docked into the density map in ChimeraX by maximizing the correlation coefficient between the experimental density map and a map generated around the model at the same resolution as the experimental density map.^65^ The voxel size and handedness of the map were checked against the model. A monomer was then extracted and manually refined in Coot.^66^ Refinements were then copied to other chains using non-crystallographic symmetry to generate a 10-mer which was then refined in PHENIX using real space refinement.^67,68^ Finally, any validation issues were manually inspected and corrected unless they had clear density to support them. Refinement and validation statistics are reported in Supplementary Table 1.

To build the atomic model of AAV2, a high resolution cryo-EM structure (PDB: 6E9D) was rigid body docked into the density map in ChimeraX by maximizing the correlation coefficient between the experimental density map and a map generated around the model at the same resolution as the experimental density map.^65^ The voxel size and handedness of the map were checked against the model. Here, the voxel size was optimized to 0.615 Å. A monomer was then extracted and manually refined in Coot.^66^ A full 60-mer capsid was generated using icosahedral symmetry matrices as implemented in ChimeraX and each monomer was assigned a unique chain ID.^65^ This 60-mer was refined in real space using PHENIX.^67,68^ The refined 60-mer was manually refined to ensure proper side chain rotamers and validate the fit of coordinated water molecules. Finally, any validation issues were manually inspected and corrected unless they had clear density to support them. Refinement and validation statistics are reported in Supplementary Table 1.

To model the AAV2-actin complex, atomic models of AAV2 and actin filaments that were determined in this study were docked into the density maps. For the full particle reconstruction of AAV2-actin, one 60-mer of AAV2 was docked into the central density (CC = 0.89) and one 16-mer of actin was docked into each of the Type A and Type B actin filament densities (Type A CC = 0.83, Type B CC= 0.81). Docked actin filaments were then copied using C3 symmetry to fill the other symmetrically related densities. For local reconstructions of Type A and Type B AAV2-actin interfaces, a 6-mer of AAV2 that corresponds to two sequential 3-fold protrusions was extracted. Then, one copy of 6-mer AAV2 and one copy of 9-mer actin filament was docked into the corresponding density (Type A – AAV2 CC = 0.92, Type A – Actin filament CC = 0.89, Type B – AAV2 CC = 0.92, Type B – Actin filament CC = 0.88).

### Tilt series data collection

AAV2-Actin grids were imaged on a Jeol CryoARM300 (RRID:SCR_024555, SCR_020163) operating at 300 kV equipped with a Gatan K3 direct electron detector, a cold field emission gun electron source, and an in-column omega energy filter with a slit width of 20 eV. First, an atlas was acquired at 50X, then medium magnification montages were collected at 4,000X and regions of interest were manually identified. 10 tilt series were then recorded with one being a region with a “low” density of AAV2-Actin bundles, six being regions with a “medium” density, and three being regions with “high” density. Tilt series were recorded using SerialEM at a nominal magnification of 15,000X, resulting in a calibrated pixel size of 2.3 Å, from -60° to 60° in 2° increments at a target underfocus of 5 μm.^59^ 1.32 e^-^/Å^2^ were used per image, resulting in a total dose of 80.8 e^-^/Å^2^ for the entire tilt series. Micrographs were not dose fractionated due to the low dose per image, instead a single integrated image was recorded at each stage tilt.

### Tomographic reconstruction and segmentation

Tilt series were aligned utilizing cross-correlation and patch tracking in IMOD.^69^ Tomograms were generated in IMOD, binned by a factor of 4 and deconvoluted with a strength of 0.7. This produced tomograms with higher contrast that were used for segmentation. A convolutional neural network in EMAN2 was trained for AAV2 capsids and actin filaments using a subset of particles from four tomograms.^70,71^ Then, the trained neural network was applied to segment all of the tomograms. Segmentations were visualized using ChimeraX.

To determine the distance between nearest neighbours, tomograms were subjected to template matching using the full particle reconstruction of the AAV2-actin complex as the search model. Doing so allowed for more accurate particle centres to be determined as compared to centres determined from image segmentation. Template matches were filtered to between 5-7σ, depending on the quality of the tomogram. Coordinates for each match were extracted from the metadata of EMAN2 and imported as atoms into ChimeraX. Nearest neighbour distance was calculated using a custom Python script, which is available upon request.

### Macromolecular visualization and figure design

Structural figures were made in ChimeraX and PyMOL 3 (Schrödinger LLC).^65^ Structures were overlayed using the matchmaker function, which utilizes a secondary structure matching algorithm. Density maps were normalized and contoured at the 0 cutoff as indicated in the figure legend.

## Supporting information

Supplementary Figure 1-11, Supplementary Table 1-3, Supplementary Video 1-4, Supplementary References

## Data Availability

The atomic model of AAV2 VLPs in KMEI was deposited in the Protein Data Bank (PDB) with PDB ID: XXXX. The corresponding density maps were also deposited in the Electron Microscopy Data Bank (EMDB) with EMDB ID: XXXXXX. The atomic model of actin filaments in KMEI was deposited in the PDB with ID: XXXX. The corresponding density maps were also deposited in the EMDB with ID: XXXXXX. The C3 full particle reconstruction of AAV2 VLPs interacting with actin filaments was deposited in the EMDB with ID: XXXXXX. The local reconstructions of AAV2 VLPs interacting with actin filaments was deposited in the EMDB with ID: XXXXXX (Type A) and XXXXXX (Type B). The 2.32 Å reconstruction of AAV2 VLPs incubated with 120 times molar excess monomeric actin was deposited in the EMDB with ID: XXXXXX. Raw micrographs were deposited in the Electron Microscopy Public Image Archive (EMPIAR) with ID: XXXXX (AAV2 VLPs alone), XXXXX (actin filaments alone), XXXXX (AAV2-actin bundles), and XXXXX (AAV2 + monomeric actin). AAV2-actin tilt series and tomograms were deposited in EMPIAR with ID: XXXXX.

## Code Availability

RELION 5 modified to support symmetry relaxation for all point groups is available at https://github.com/LSB-Helsinki/relion5/tree/relax.

## Acknowledgements

This work was supported by National Institute of General Medical Sciences Grant R01GM082946 to R.M. and National Institute of Deafness and Other Communication Disorders Grant R01DC018827 to J.B. We thank other members of the McKenna and Bird labs for helpful comments and discussions during the preparation of this manuscript. We thank the University of Florida ICBR Electron Microscopy Core Facility (RRID:SCR_019146) for access to negative staining facilities and are grateful to Mary Gragg and Paul Chipman for expert assistance. We thank the Herbert Wertheim UF Scripps Institute for Biomedical Innovation & Technology – Structural Biology Cryo-Electron Microscopy Core (RRID:SCR_024555) for access to cryo-EM and cryo-ET data collection using the JEOL Cryo ARM 300 (RRID:SCR_020163) and Leica GP2 plunge freezer, which was essential to collecting AAV2-actin data sets. The authors wish to acknowledge CSC – IT Center for Science, Finland, for computational resources. M.G. would like to thank Linda Bloom and members of the Bloom laboratory for helpful comments and discussions during the preparation of this manuscript.

## Ethics declarations

## Contributions

M.G. contributed to all aspect of this study, experimental design, sample production, cryo-EM, data processing, data interpretation, and writing of the manuscript. J.H. and J. B. contributed to actin production, and all actin associated binding and fluorescence microscopy studies. A.S.-G. cryo-EM and tomography data collection. M.P. and J.H-B. AAV production, and negative stain EM. T.H. confocal microscopy studies. J.T.H. in code writing and data processing of the cryo-EM AAV2-actin complex. X.S. provided validation of computational methods. A.B., M.M. and J.B. in discussion and manuscript preparation. R.M. with M.G. contributed to the concept of the study and writing the first manuscript draft. All authors contributed to writing and approving the final version of the manuscript.

