## Supplementary Figure 1-11, Supplementary Table 1-3, Supplementary Video 1-4, Supplementary References for "AAV2 Crosslinks Actin Filaments: Implications for AAV Gene Therapy Vector Design"

Supplementary Table 1-3

Supplementary Video 1-4

Supplementary References

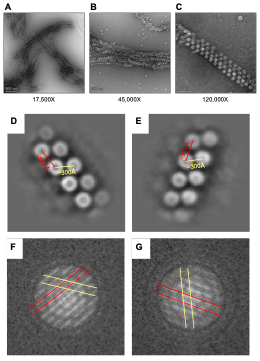

**Supplementary Figure 1. AAV2 bundles actin filaments with high periodicity.** (A-C) Negative stain electron micrographs of AAV2 interacting with actin filaments at 17,500X magnification (A), 45,000X magnification (B), or 120,000X magnification (C). Scales of images are denoted by the scale bar in the lower left corner, and correspond to 500 nm (A), 200 nM (B), and 100 nm (C). (D-E) 2D class averages using a box size of 256 pixels (2109 Å). Estimated distances along the bundle and between bundles are shown in red and yellow, respectively. (F-G) Fourier transform of the 2D class average shown in D-E, respectively. Exemplary layer lines corresponding to the distances shown in D-E are highlighted with red and yellow lines, colored as in D-E.

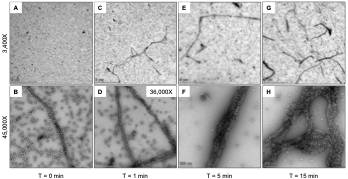

**Supplementary Figure 2. AAV2 bundles actin filaments cooperatively.** (A-H) Negative stain electron micrographs at 3,400X magnification (A, C, E, G), 36,000X magnification (D), and 45,000X magnification (B, F, H). Samples were incubated for 0 minutes (A-B), 1 minute (C-D), 5 minutes (E-F), and 15 minutes (G-H) before being applied to the grid and given 1 minute to adsorb to the carbon surface. Scale bars are indicated in the lower left corner.

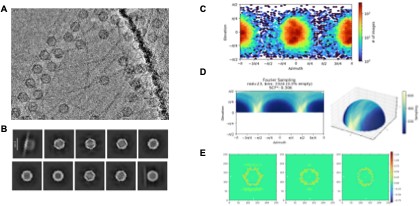

**Supplementary Figure 3. AAV2 exhibits strong preferred orientation when bound to actin filaments.** (A) Cryogenic electron micrograph of AAV2 bound to actin filaments. Clear periodicity is readily observed. (B) 2D class averages from early in cryo-EM data processing. Actin filaments can be seen in all classes, but they are less defined than the capsid density. (C) Angular distribution plot from midway through cryo-EM data processing. Two orientations of AAV2 that are rotated 180° from each other are heavily preferred. (D) Fourier sampling plot showing significant preferred orientation. Note that the corrected sampling compensation factor (SCF*) is 0.306. Isotropic maps have a SCF* of 0.81 or greater. (E) Three orthogonal real space slices through a 3D reconstruction midway through cryo-EM data processing. Densities corresponding to actin filaments are clearly seen, but have much lower intensity than the capsid density, likely due to significant map anisotropy.

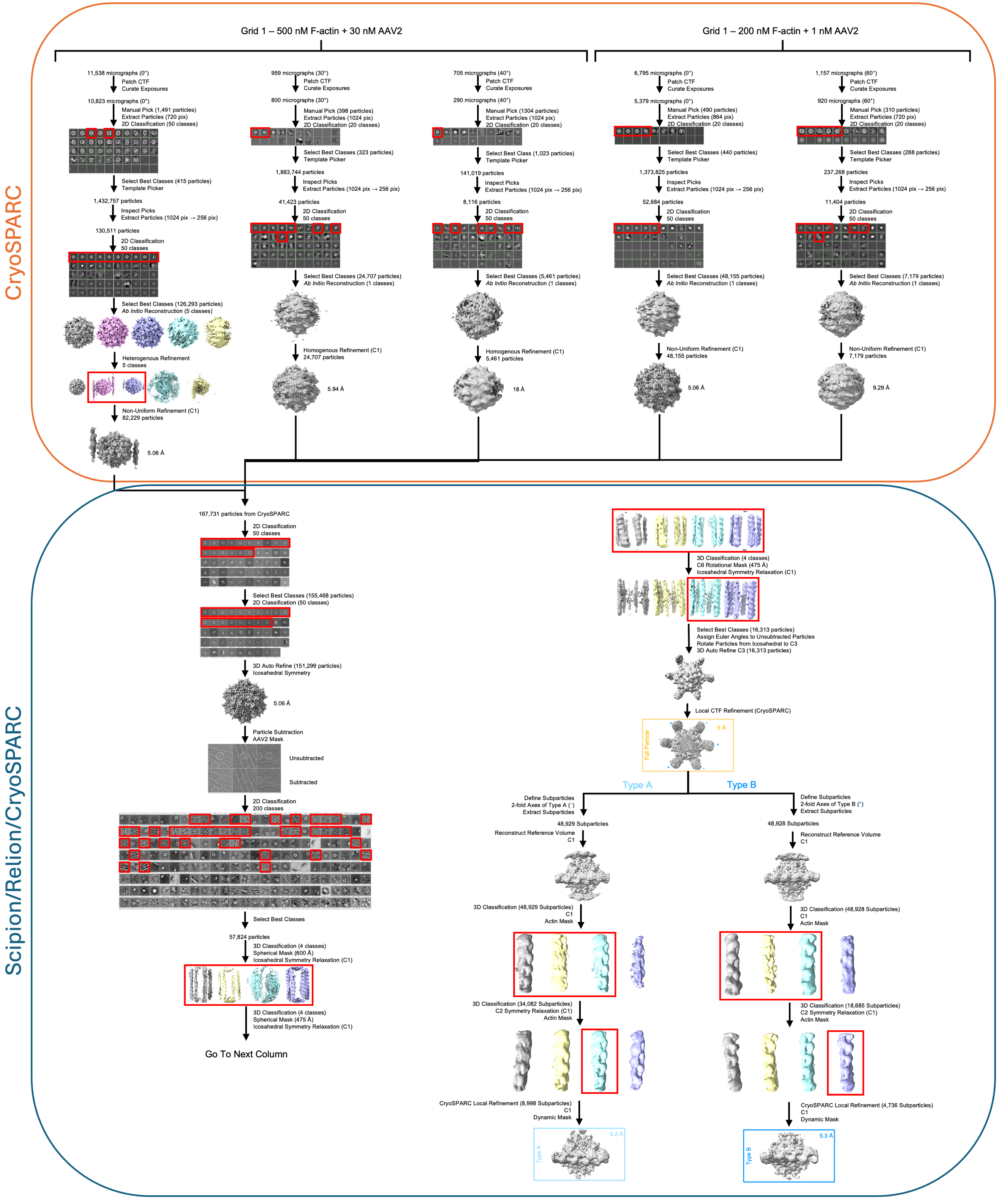

**Supplementary Figure 4. Cryo-EM workflow for reconstructing AAV2 bound to actin filaments.** Data processing workflow that was used to produce the three reconstructions used in this work. Initial particle processing was conducted in CryoSPARC v4.7.1. Downstream processing was conducted in Scipion3 using RELION5, Xmipp, and CryoSPARC plugins. The 5.0 Å full particle reconstruction is indicated by the orange box, the 5.2 Å local reconstruction of the Type A AAV2-actin interface is indicated by the light blue box, and the 5.3 Å local reconstruction of the Type B AAV2-actin interface is indicated by the dark blue box. Selected classes that were used in further steps indicated by the red boxes.

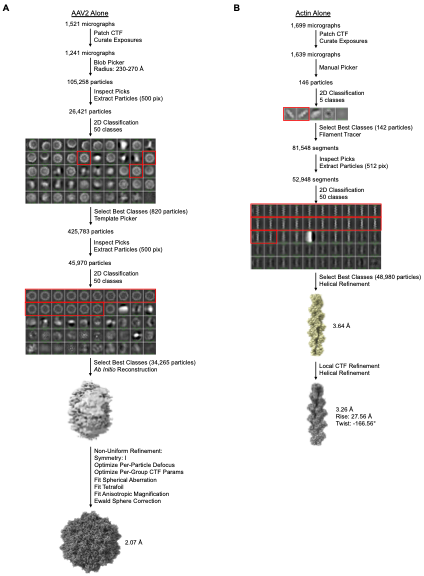

**Supplementary Figure 5. Cryo-EM workflow for reconstructing AAV2 and actin filaments alone.** (A) Cryo-EM workflow used to determine the 2.07 Å reconstruction of AAV2 VLPs in KMEI. Selected classes are shown in red boxes. (B) Cryo-EM workflow used to determine the 3.26 Å reconstruction of actin filaments in KMEI. Selected classes are shown in red boxes.

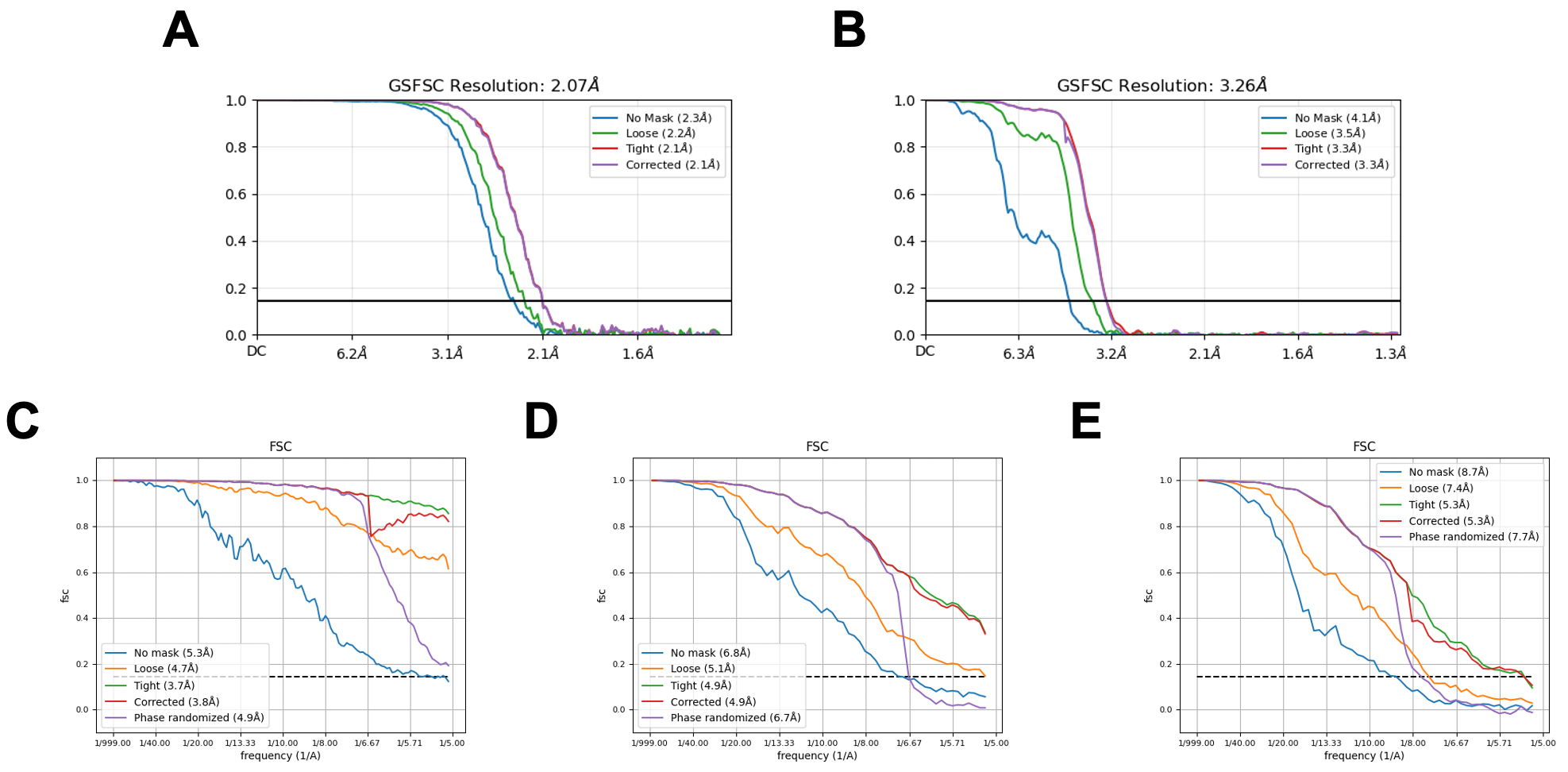
**Supplementary Figure 6. Validation metrics from cryo-EM reconstructions.** (A-E) Gold standard Fourier shell correlation (GSFSC) plots used to determine the resolution of the reconstruction for AAV2 alone (A), actin filaments alone (B), AAV2-actin C3 full particle (C), AAV2-actin Type A local (D), and AAV2-actin Type B local (E).

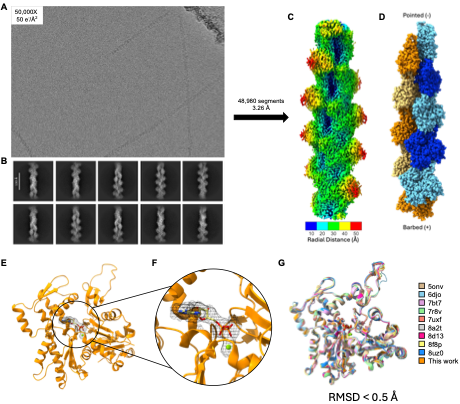

**Supplementary Figure 7. Atomic resolution reconstruction of actin filaments in KMEI.** (A) Cryogenic electron micrograph of single actin filaments in KMEI buffer. (B) 2D class averages showing well defined features and secondary structure. (C) 3.26 Å electron density map generated by helical reconstruction. The surface is colored radially according to the color code below. (D) Atomic model of 10-mer actin that was refined using the electron density map. Each strand of the actin double helix is colored either orange or blue, with subunits on the same strand alternating between a light and dark shade. The barbed and pointed ends are indicated. (E) Cartoon representation of the refined actin monomer. The bound ADP is shown in yellow stick form alongside the bound Mg^2+^ ion in green. The electron density map for the ADP is shown contoured at 1.5 σ. (F) Same as in E, but zoomed to show more detail of the ADP-Mg^2+^ binding site. (G) Overlay between the atomic model of actin monomer determined in this study (shown in orange) and nine previously determined structures of actin (colored as indicated in the figure). The root mean square deviation was below 0.5 Å across all Cα atoms in the atomic model.

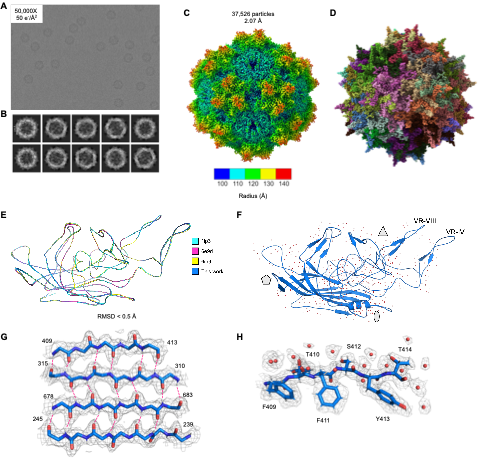

**Supplementary Figure 8. Atomic resolution reconstruction of AAV2 VLPs in KMEI.** (A) Cryogenic electron micrograph of AAV2 VLPs alone in KMEI buffer. (B) 2D class averages showing multiple particle orientations, strong features, and secondary structure. (C) 2.07 Å electron density map with the surface colored radially from the center, as indicated by the color bar below. (D) Atomic model of AAV2 refined based on the electron density map. Note that the capsid is an icosahedral 60-mer of the monomeric viral capsid protein. Water molecules are shown as red spheres. (E) Overlay of the AAV2 monomer determined in this work (dark blue) with three previously determined structures (colored as indicated in the figure). The root mean square deviation was below 0.5 Å across all Cα atoms in the atomic model. (F) Atomic model of AAV2 determined here shown in cartoon representation, with secondary structure highlighted. The 468 spherical densities that were modeled as water molecules are shown as red spheres. The grey pentagon represents the icosahedral 5-fold symmetry axis, the grey triangle represents the icosahedral 3-fold symmetry axis, and the grey oval represents the icosahedral 2-fold symmetry axis. VR-IV and VR-VIII, which are important for actin binding, are labeled. (G) A section of four anti-parallel β sheets found in the core of the jelly roll motif of AAV2 shown in stick representation. Start and end residues are labeled. The electron density is shown as a mesh contoured at 1.5 σ. Backbone hydrogen bonds are shown as dashed pink lines. Electronic overlap between hydrogen bonds can be directly visualized. (H) A heavily hydrated section of AAV2 is shown with protein atoms in stick form and water molecules as red spheres. The identity of the residues are labeled. The electron density is shown as a mesh contoured at 1.5 σ.

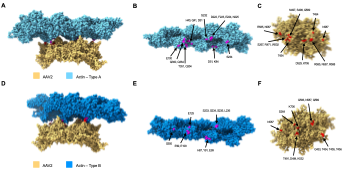

**Supplementary Figure 9. Residues that mediate the interaction between AAV2 and actin.** (A) 6-mer of AAV2 (orange) and 9-mer of actin (light blue) as determined by docking into the local reconstruction of the Type A interface. AAV2 residues that are within 5 Å of actin are colored red. Actin residues that are within 5 Å of AAV2 are colored magenta. (B) 9-mer model of actin with AAV2 contact residues colored magenta and labeled. (C) 6-mer model of AAV2 with actin contact residues colored red and labeled. (D) 6-mer of AAV2 (orange) and 9-mer of actin (dark blue) as determined by docking into the local reconstruction of the Type B interface. AAV2 residues that are within 5 Å of actin are colored red. Actin residues that are within 5 Å of AAV2 are colored magenta. (E) 9-mer model of actin with AAV2 contact residues colored magenta and labeled. (F) 6-mer model of AAV2 with actin contact residues colored red and labeled.

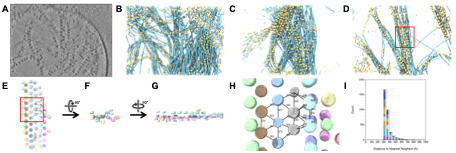

**Supplementary Figure 10. Cryo-electron tomograms reveal a hexagonal lattice of AAV2 capsids among heterogeneous actin filaments.** (A) Central slice through an AAV2-actin tomogram. (B) Segmented tomogram where volumes corresponding to AAV2 capsids are colored orange and volumes corresponding to actin filaments are colored light blue. Dense, heterogeneous bundles are visualized. (C-D) Two other examples of AAV2-actin segmented tomogram, colored as in B. The red box in D indicates the area zoomed in on in E. (E) Segmented tomographic volumes corresponding to a hexagonal lattice of AAV2 capsid along actin filaments. Capsid volumes are colored according to the co-linear bundle they are in. The hexagonal lattice is shown by black connecting lines for a subset of capsid volumes. The red box indicates the area zoomed in on in H. (F) Same as in E, but rotated 90° around the x axis. (G) Same as in F, but rotated 90° around the y axis. (H) Close up view of the hexagonal lattice of AAV2. Distances between volume centers (in Å) found by template matching are labeled. Note the uniformity of distances. (I) Histogram showing the nearest neighbor distance between AAV2 particle centers found by template matching across ten tomograms. Each tomogram is colored separately. Note that the vast majority of nearest neighbor distances are between 300-400 Å.

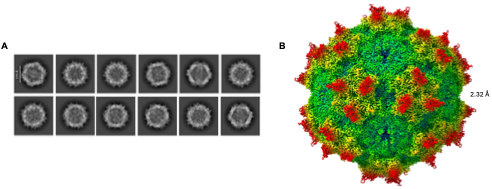

**Supplementary Figure 11. AAV2 does not bind monomeric actin.** (A-B) 2D class averages (A) and 2.32 Å reconstruction (B) from AAV2 VLPs incubated with 120 times molar excess monomeric actin. No density is visible that is not attributable to the AAV2 capsid.

**Supplementary Table 1. Cryo-EM data collection, refinement and validation statistics.**

|  | AAV2 Alone  (EMDB-xxxx)  (PDB xxxx) | Actin Alone  (EMDB-xxxx)  (PDB xxxx) | AAV2-Actin (Full Particle)  (EMDB-xxxx) | AAV2-Actin (Type A Interface)  (EMDB-xxxx) | AAV2-Actin (Type B Interface)  (EMDB-xxxx) |
| --- | --- | --- | --- | --- | --- |
| **Data collection and processing** |  |  |  |  |  |
| Magnification | 50,000 | 50,000 | 50,000 | 50,000 | 50,000 |
| Voltage (kV) | 300 | 300 | 300 | 300 | 300 |
| Electron exposure (e–/Å^2^) | 50 | 50 | 50 | 50 | 50 |
| Defocus range (μm) | 0.8-1.8 | 0.8-1.8 | 0.8-1.8 | 0.8-1.8 | 0.8-1.8 |
| Pixel size (Å) | 0.62 | 0.62 | 0.62 | 0.62 | 0.62 |
| Symmetry imposed | I | Helical | C3 | C1 | C1 |
| Initial particle images (no.) | 45,970 | 81,458 | 167,731 | 48,929 | 48,928 |
| Final particle images (no.) | 34,265 | 48,980 | 16,313 | 8,998 | 4,736 |
| Map resolution (Å)  FSC threshold | 2.07  0.143 | 3.26  0.143 | 5.0  0.143 | 5.2  0.143 | 5.3  0.143 |
| **Refinement** |  |  |  |  |  |
| Initial model used (PDB code) | 6E9D | 8A2T |  |  |  |
| Model resolution (Å)  FSC threshold | 2.14  0.5 | 3.36  0.5 |  |  |  |
| Masked Correlation Coefficient | 0.86 | 0.89 |  |  |  |
| Map sharpening *B* factor (Å^2^) | -60.9 | -86.7 |  |  |  |
| Model composition (per monomer)  Non-hydrogen atoms  Protein residues  Ligand atoms  Water | 4319  520  0  468 | 2911  371  28  0 |  |  |  |
| *B* factors (Å^2^)  Protein  Ligand  Water | 18.07  N/A  20.39 | 47.11  36.52  N/A |  |  |  |
| R.m.s. deviations  Bond lengths (Å)  Bond angles (°) | 0.011  1.133 | 0.009  0.854 |  |  |  |
| Validation  MolProbity score  Clashscore  Poor rotamers (%) | 1.92  11.53  0.7 | 2.28  8.77  0.0 |  |  |  |
| Ramachandran plot  Favored (%)  Allowed (%)  Disallowed (%) | 97.04  2.96  0 | 93.24  6.76  0 |  |  |  |

**Supplementary Table 2. Residues on the surface of AAV2 that contact actin, heparin, and AAVR.**

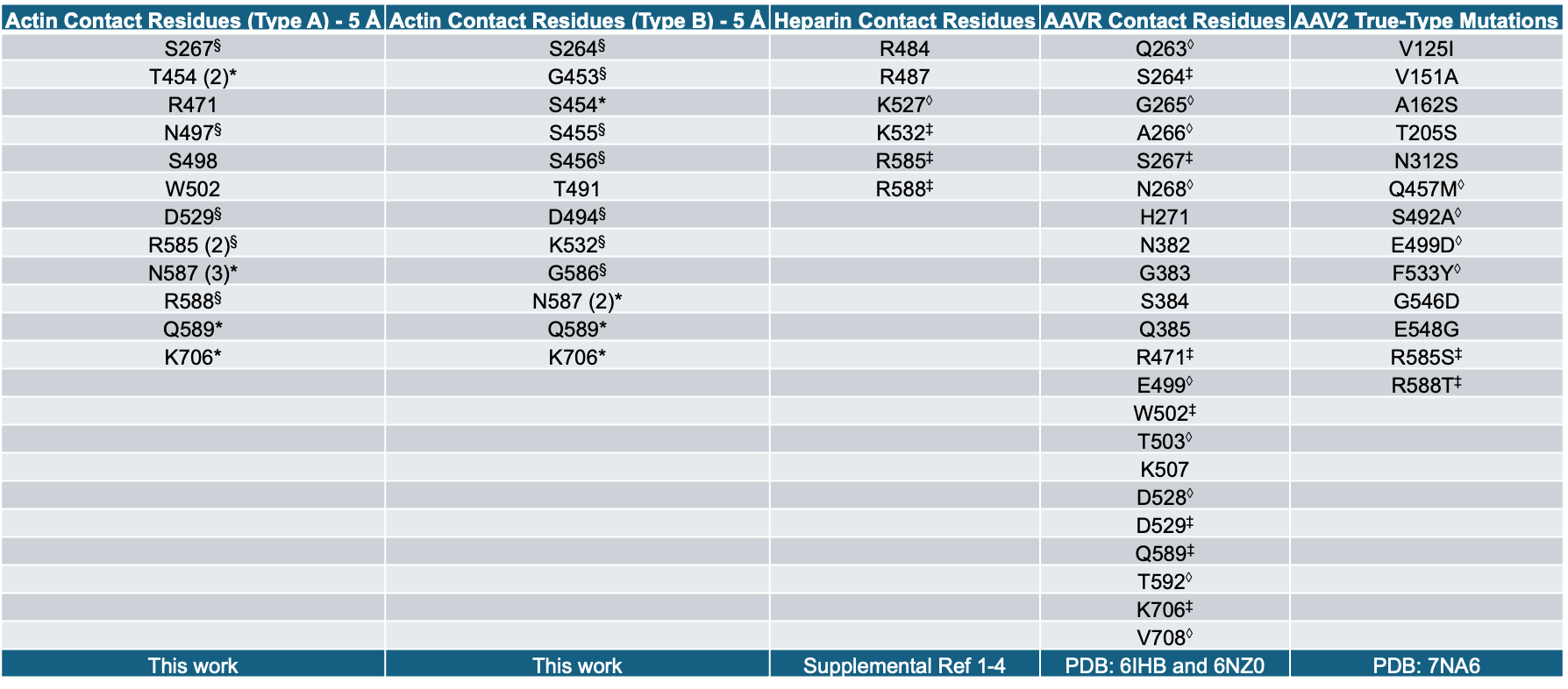

* denotes contact residues that are shared between Type A actin binding and Type B actin binding. § denotes contact residues that are within three residues between Type A actin binding and Type B actin binding. ‡ denotes residues that are shared between at least one binding mode of actin and heparin, AAVR, or are mutated in AAV2 true-type. ◊ denotes residues that are within three residues of at least one binding mode of actin and heparin, AAVR, or are mutated in AAV2 true-type.

**Supplementary Table 3. Residues on the surface of actin that contact AAV2 and other actin binding proteins.**

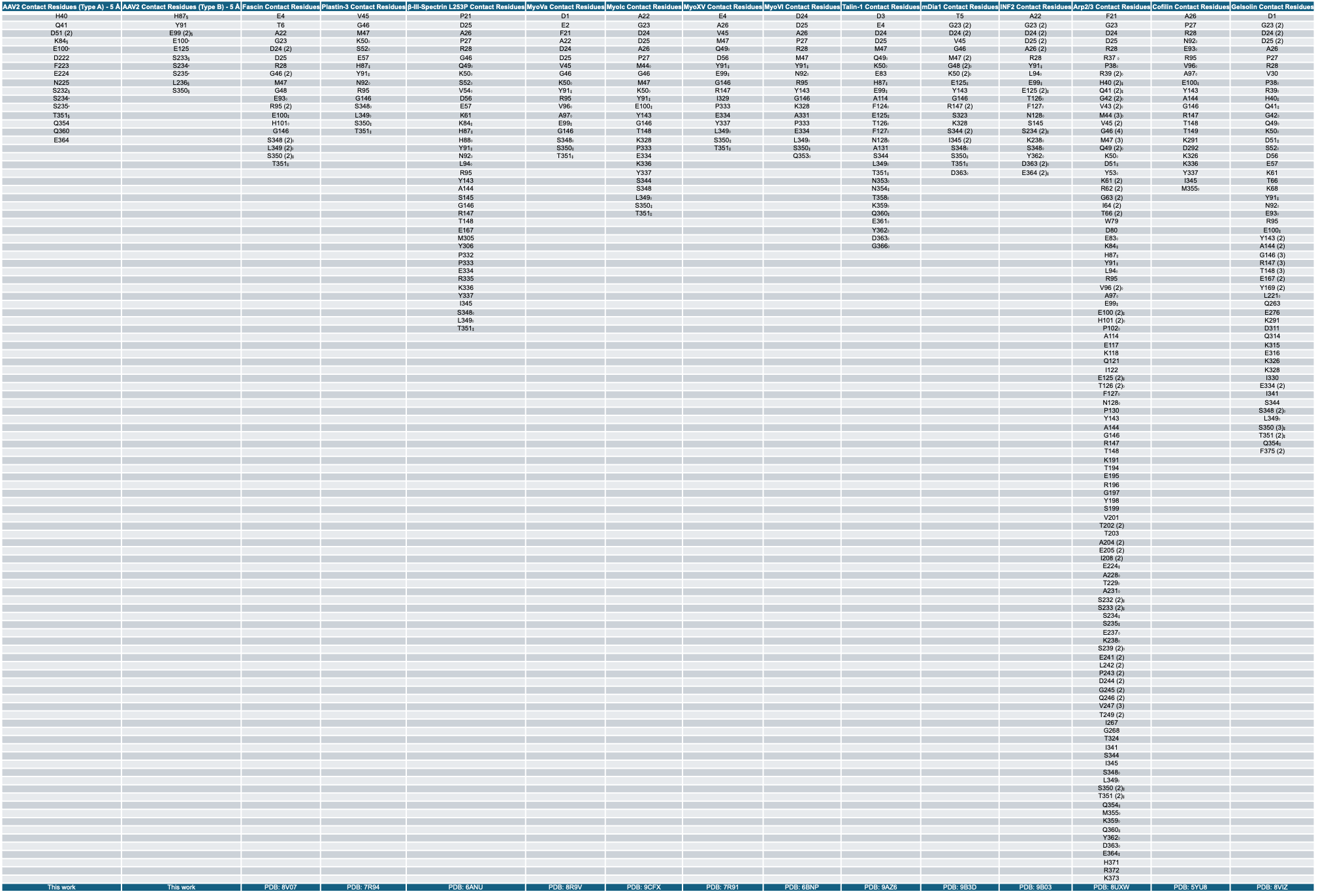

* denotes contact residues that are shared between Type A AAV2 binding and Type B AAV2 binding. § denotes contact residues that are within three residues between Type A AAV2 binding and Type B AAV2 binding. ‡ denotes residues that are shared between at least one binding mode of AAV2 and other actin binding proteins. ◊ denotes residues that are within three residues of at least one binding mode of AAV2 and other actin binding proteins.

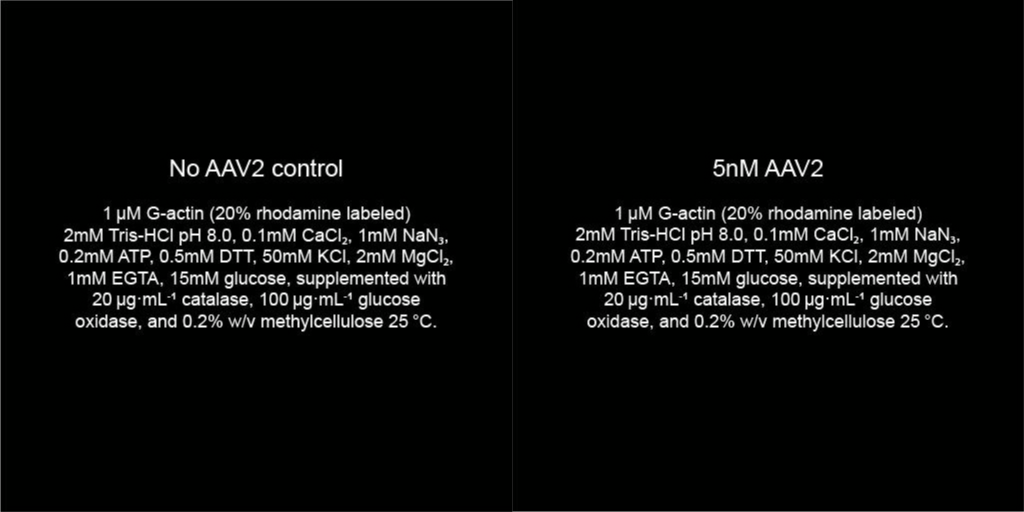

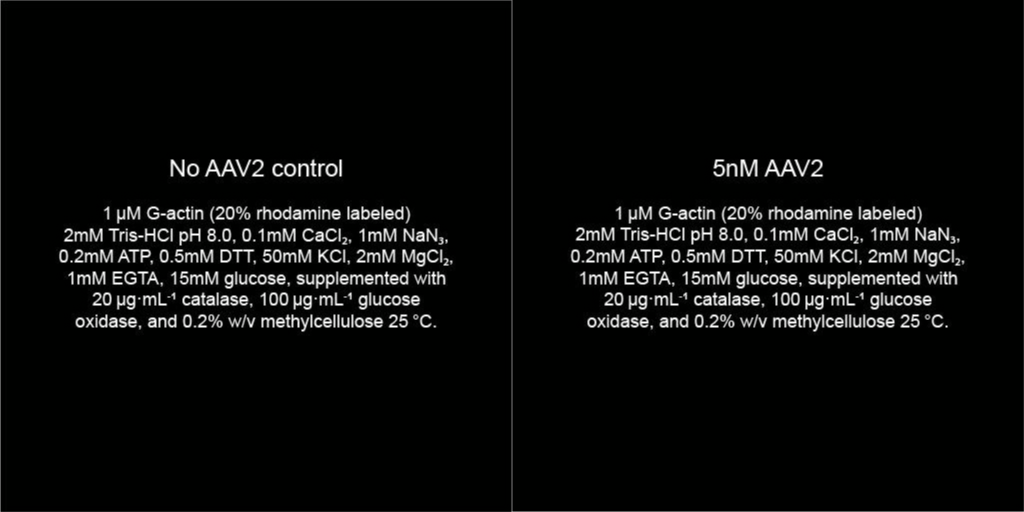
**Supplementary Video 1. Cropped TIRFM video of actin alone versus actin with AAV2 undergoing polymerization.**

**Supplementary Video 2. Full screen TIRFM video of actin alone versus actin with AAV2 undergoing polymerization.**

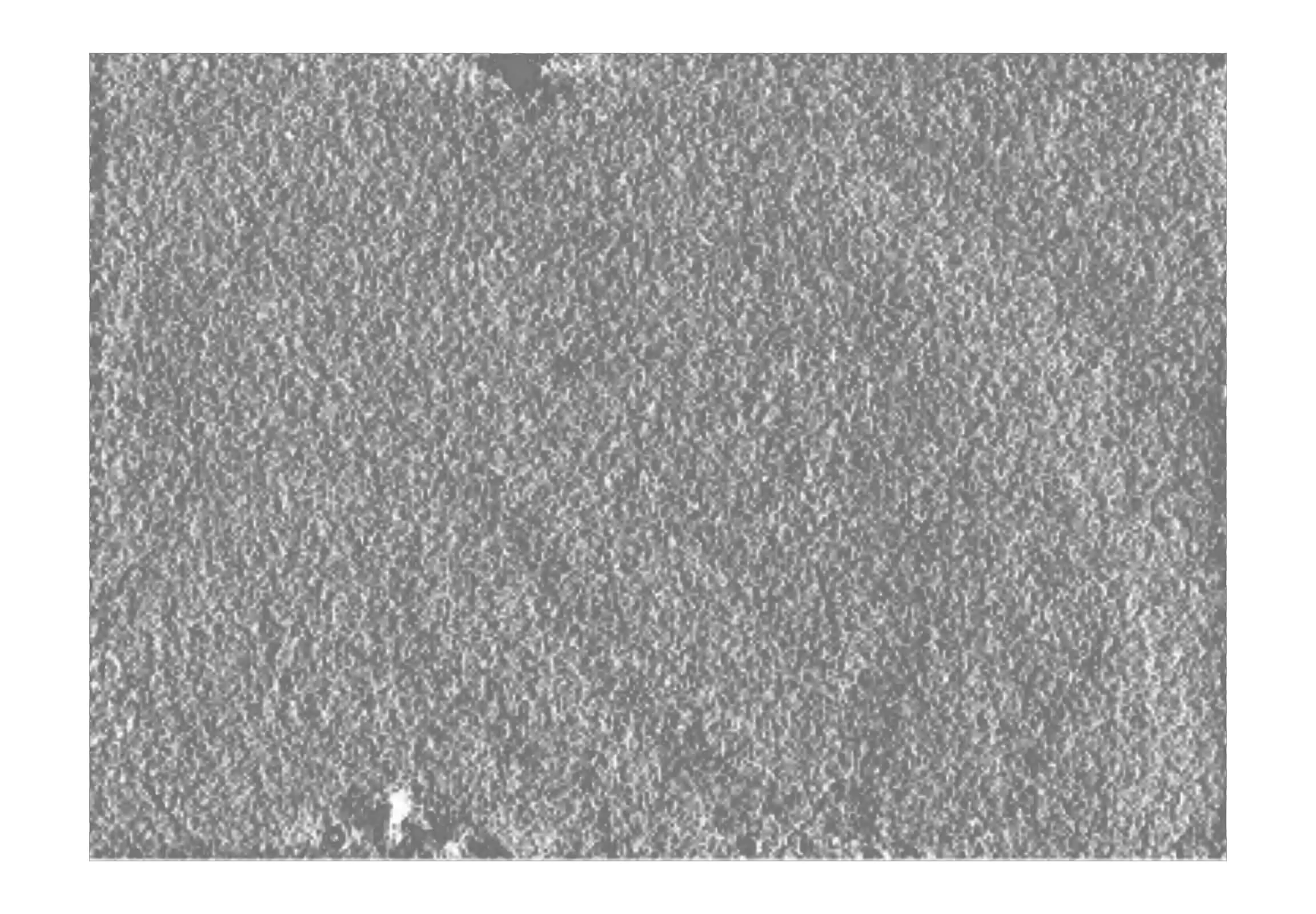

**Supplementary Video 4. Z-stack video of an AAV2-actin tomogram.**

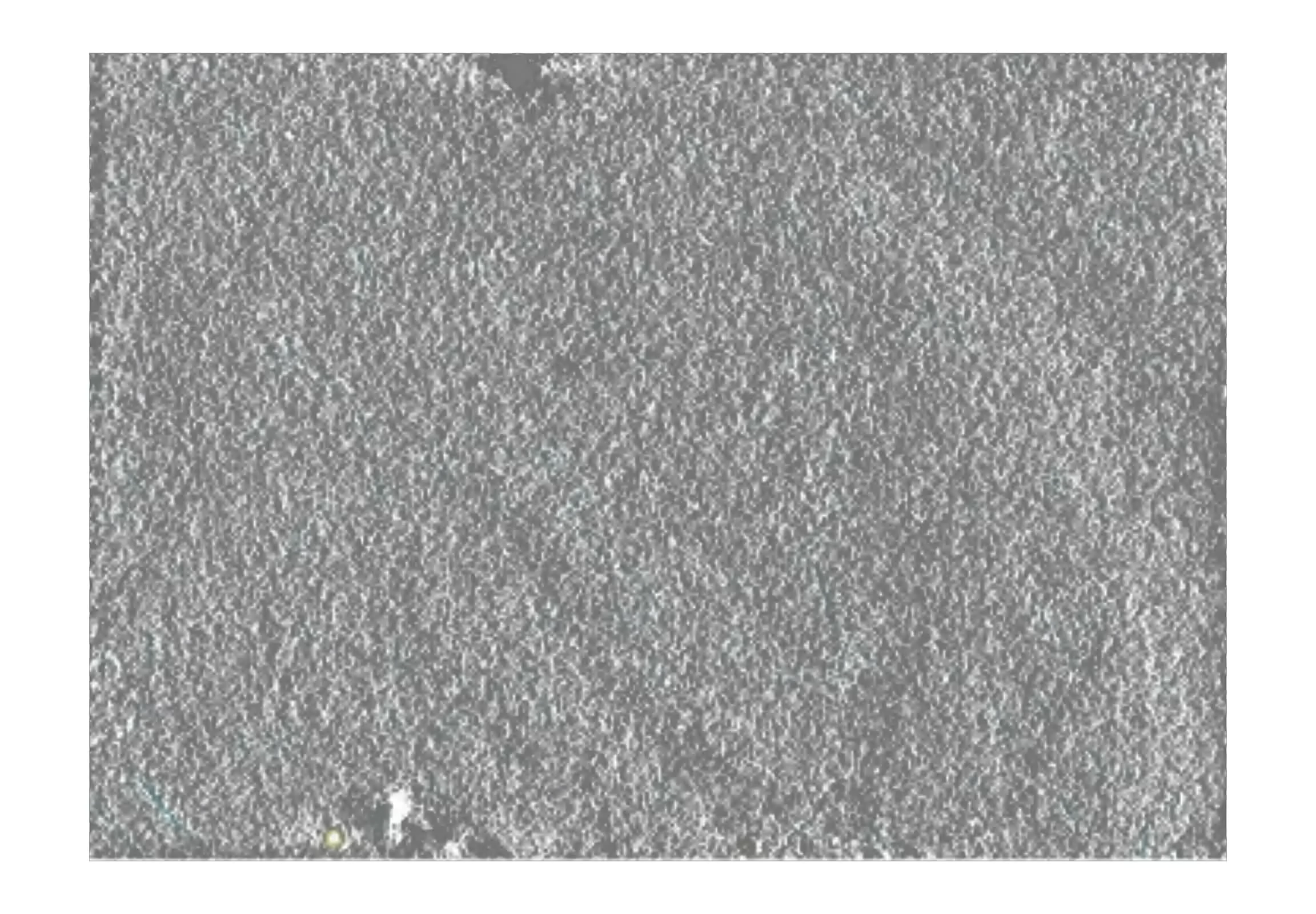

**Supplementary Video 5. Segmented Z-stack video of an AAV2-actin tomogram.** Densities that were segmented as capsid are colored yellow and densities that were segmented as actin filaments are colored light blue.

**Supplemental References:** ^1–4^

1. O’Donnell, J., Taylor, K. A. & Chapman, M. S. Adeno-associated virus-2 and its primary cellular receptor—Cryo-EM structure of a heparin complex. *Virology* **385**, 434–443 (2009).

2. Kern, A. *et al.* Identification of a Heparin-Binding Motif on Adeno-Associated Virus Type 2 Capsids. *J. Virol.* **77**, 11072–11081 (2003).

3. Opie, S. R., Warrington, K. H., Agbandje-McKenna, M., Zolotukhin, S. & Muzyczka, N. Identification of Amino Acid Residues in the Capsid Proteins of Adeno-Associated Virus Type 2 That Contribute to Heparan Sulfate Proteoglycan Binding. *J. Virol.* **77**, 6995–7006 (2003).

4. Bennett, A. *et al.* Comparative structural, biophysical, and receptor binding study of true type and wild type AAV2. *J. Struct. Biol.* **213**, 107795 (2021).
